# The *Toxoplasma gondii* homolog of ATPase inhibitory factor 1 is critical for mitochondrial cristae maintenance and stress response

**DOI:** 10.1101/2024.08.09.607411

**Authors:** Madelaine M. Usey, Anthony A. Ruberto, Diego Huet

**Affiliations:** Department of Cellular Biology, University of Georgia, Athens, GA, USA; Department of Pharmaceutical and Biomedical Sciences, University of Georgia, Athens, GA, USA; Center for Tropical and Emerging Global Diseases, University of Georgia, Athens, GA, USA; Institute of Bioinformatics, University of Georgia, Athens, GA, USA

## Abstract

The production of energy in the form of ATP by the mitochondrial ATP synthase must be tightly controlled. One well-conserved form of regulation is mediated via ATPase inhibitory factor 1 (IF1), which governs ATP synthase activity and gene expression patterns through a cytoprotective process known as mitohormesis. In apicomplexans, the processes regulating ATP synthase activity are not fully elucidated. Using the model apicomplexan *Toxoplasma gondii*, we found that knockout and overexpression of TgIF1, the structural homolog of IF1, significantly affected gene expression. Additionally, TgIF1 overexpression resulted in the formation of a stable TgIF1 oligomer that increased the presence of higher order ATP synthase oligomers. We also show that parasites lacking TgIF1 exhibit reduced mitochondrial cristae density, and that while TgIF1 levels do not affect growth in conventional culture conditions, they are crucial for parasite survival under hypoxia. Interestingly, TgIF1 overexpression enhances recovery from oxidative stress, suggesting a mitohormetic function. In summary, while TgIF1 does not appear to play a role in metabolic regulation under conventional growth conditions, our work highlights its importance for adapting to stressors faced by *T. gondii* and other apicomplexans throughout their intricate life cycles.

**SIGNIFICANCE STATEMENT:**

- *Toxoplasma gondii* is a member of the Apicomplexa, a phylum consisting of parasites responsible for significant global morbidity and mortality. An intact mitochondrial ATP synthase is critical *T. gondii* survival, but how this enzyme is regulated is not completely understood.
- Our work demonstrates that the *T. gondii* homolog of ATPase inhibitory factor 1 (TgIF1) does not impact metabolism under standard culture conditions, but plays a role in mitochondrial cristae density and stress responses.
- This study reveals the role of TgIF1 in regulating ATP synthase activity under stressful conditions and increases our understanding of this divergent enzyme in *T. gondii*.

## INTRODUCTION

As the power generator within the mitochondrion, the primary function of the ATP synthase is to generate energy in the form of ATP. Although a high energy output from the ATP synthase is often necessary for cellular survival, this output must be tightly regulated so that cells can adapt to environmental changes, stressors, and varying metabolic demands. One key interactor protein involved in ATP synthase regulation is ATPase inhibitory factor 1 (IF1) (Pullman and Monroy, 1963). IF1 binds to the F_1_ portion of the enzyme, between the α and the catalytic β subunits (Cabezón *et al*., 2003). The binding of IF1 at this interface disrupts the cyclical conformational changes in the catalytic β subunit that are necessary for ATP production, thus locking the enzyme in an inactive state (Gledhill *et al*., 2007; Jonckheere *et al*., 2012).

IF1 was initially identified as an ATP synthase inhibitor in bovine mitochondria and has since been studied extensively in plants, mammals, and yeast (Pullman and Monroy, 1963; Cintrón and Pedersen, 1979; Hashimoto *et al*., 1981; Norling *et al*., 1990). As IF1 activity has been shown to be regulated by pH, it was initially thought that IF1 only inhibited the reverse, or hydrolytic, function of the ATP synthase, particularly under hypoxic conditions that result in mitochondrial matrix acidification (Cabezon *et al*., 2000; Gore *et al*., 2022). However, various studies have shown that IF1 is also capable of inhibiting the synthetic activity of the enzyme (Zanotti *et al*., 2009; Sanchez-Cenizo *et al*., 2010; Formentini *et al*., 2014, 2017; Santacatterina *et al*., 2016; Kahancová *et al*., 2020; Esparza-Moltó *et al*., 2021).

Interestingly, IF1 inhibition of the ATP synthase under normoxic conditions can induce mitohormesis, a process involving the activation of cell signaling pathways to support cellular health in response to stress (Yun and Finkel, 2014). This activation occurs as IF1 binding blocks the backflow of protons into the mitochondrial matrix, resulting in membrane potential hyperpolarization and the production of mitochondrial reactive oxygen species (mtROS). Acting as retrograde signaling molecules, mtROS can travel to the nucleus and modulate gene expression, promoting the activation of pathways involved in cell survival, repair, and antioxidant defense. As a result, the cell is then better prepared to handle future mitochondrial stressors (Esparza-Molto *et al*., 2017; García-Aguilar and Cuezva, 2018).

Although the role of IF1 has been investigated extensively in a wide range of organisms (Pullman and Monroy, 1963; Cintrón and Pedersen, 1979; Hashimoto *et al*., 1981; Norling *et al*., 1990), its characterization in protozoan parasites remains largely unexplored. Protozoan parasites include the causative agents of diseases such as malaria, toxoplasmosis, trypanosomiasis, and cryptosporidiosis. These diseases are a major burden on global health, resulting in over a million deaths each year and severe socioeconomic consequences (Ung *et al*., 2021). As many of the drugs for treating infections caused by these parasites have become less effective (Rao *et al*., 2023), it is critical that we develop a better understanding of pathways essential for parasite viability. This will facilitate the development of novel therapeutic approaches to prevent and treat protozoan-borne diseases. Because the mitochondrion plays a critical role in energy production and cellular health of many of these parasites, elucidating its regulatory mechanisms, such as those involving IF1, could support novel therapeutic strategies.

The protozoan parasite *Toxoplasma gondii*, which causes toxoplasmosis, possesses a ortholog of IF1 (TgIF1), although its function has not yet been characterized. TgIF1 (TGME49_215350) was initially discovered associated with the *T. gondii* ATP synthase through immunoprecipitation, and its presence in the enzymatic complex was later confirmed via cryo-electron microscopy and complexome studies (Huet *et al*., 2018; Maclean *et al*., 2021; Muhleip *et al*., 2021). Intriguingly, TgIF1 is not conserved on the amino acid sequence level when compared to yeast and mammalian IF1. Instead, it was originally identified as a putative IF1 ortholog using secondary structure prediction algorithms (Huet *et al*., 2018; Zimmermann *et al*., 2018). Additionally, TgIF1 appears to be conserved in the apicomplexan *Plasmodium falciparum* and it is not conserved in *Cryptosporidium spp.*, another apicomplexan genus with highly reduced mitochondria (Huet *et al*., 2018; Tsaousis and Keithly, 2019). Nonetheless, it remains unclear whether TgIF1 performs similar roles to IF1 found in other organisms. Therefore, we aimed to characterize the role of TgIF1 in *T. gondii*. To do so, we utilized CRISPR/Cas9 to create stable TgIF1 knockout and overexpression lines. We then we used a combination of genome-wide profiling, biochemical techniques, microscopy methods, and phenotypic experiments to understand the function of TgIF1 within the ATP synthase complex, its impact on metabolic flux and mitochondrial morphology, and its role in the parasite’s stress response. Our work revealed that although TgIF1 does not impact *T. gondii* metabolism under conventional growth conditions, it plays a key role in mitochondrial cristae maintenance as well as in the parasite’s ability to respond to oxidative and hypoxic stress.

## RESULTS

### Generation and phenotypic characterization of IF1^Ty^, IF1^KO^, and IF1^Over^ strains

To begin the characterization of TgIF1, we first utilized CRISPR/Cas9 and homology-directed repair to tag the C terminus of TgIF1 with an in-frame Ty epitope tag, thus creating the IF1^Ty^ strain (Figure 1A). We tagged the C terminus of the protein because structural studies in mammals have confirmed that the N terminal region of IF1 is critical for the interaction of the inhibitor with the ATP synthase (Gledhill *et al*., 2007). We then replaced the entire Ty-tagged TgIF1 gene with a pyrimethamine-resistant dihydrofolate-reductase (DHFR) cassette using homology directed repair (Figure 1B). The generation of a complete IF1 knockout (IF1^KO^) strain was possible because TgIF1 was previously predicted to be non-essential for the lytic cycle of *T. gondii* (Sidik et al., 2016). Finally, the IF1^KO^ strain was used as the genetic background for overexpression of TgIF1 (IF1^Over^). To generate this strain, an exogenous copy of TgIF1 with a C-terminal HA tag under the control of a strong Tub8 promoter was targeted to the parasite *UPRT* locus (Figure 1C).

**Figure 1.**
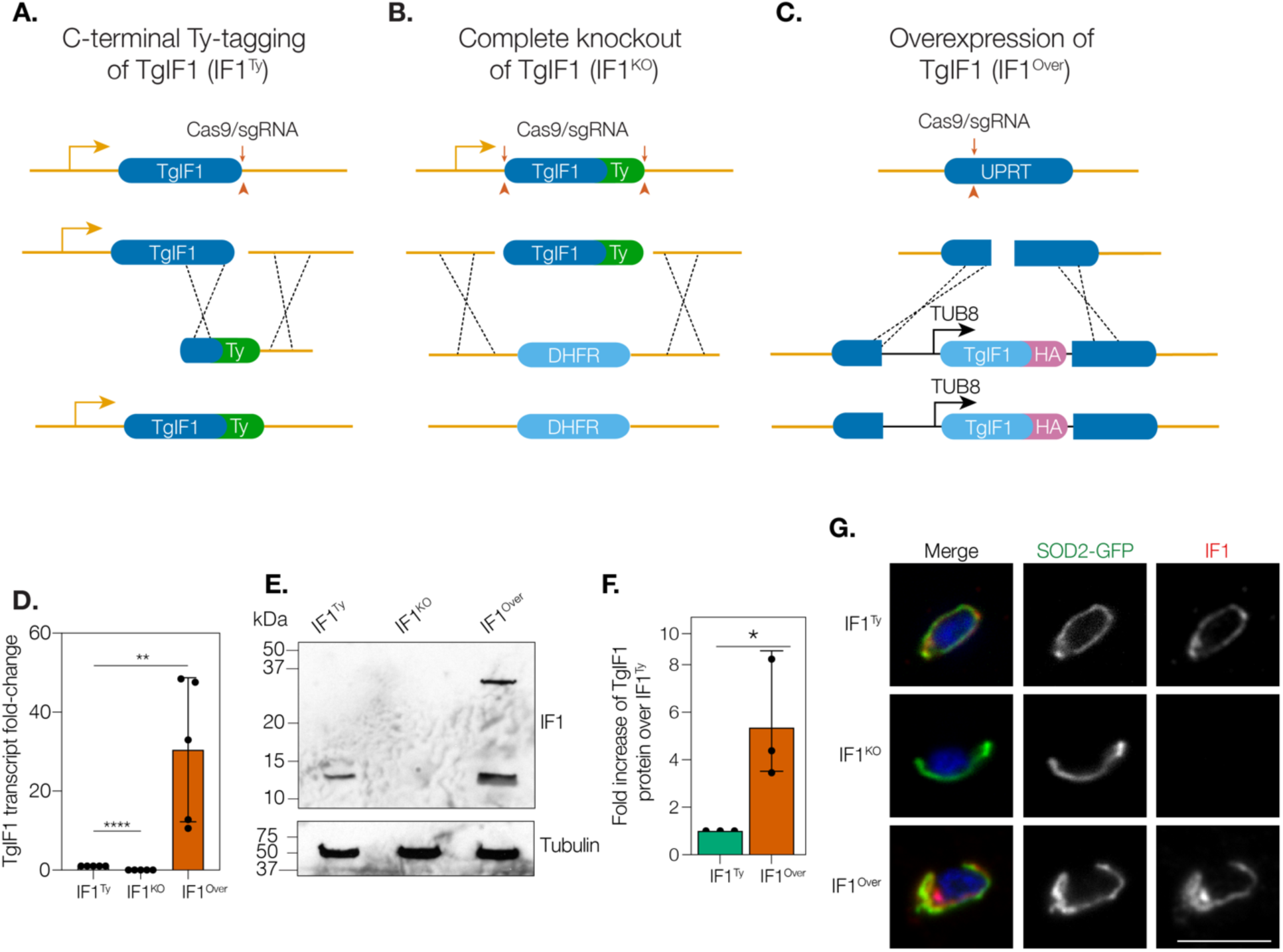
Generation of the IF1^Ty^, IF1^KO^ and IF1^Over^ strains. **A.** Schematic representation of the strategy used to generate the IF1^Ty^ strain. **B.** To create a complete knockout of TgIF1 (IF1^KO^), the TgIF1 locus in the IF1^Ty^ strain was replaced with a dihydrofolate reductase (DHFR) cassette using CRISPR/Cas9 homology-directed repair. **C.** Overexpression of TgIF1 was achieved by the exogenous expression of an HA-tagged TgIF1 copy driven by the strong Tub8 promoter from the uracil phosphoribosyltransferase (UPRT) (TGME49_312480) locus. **D.** Quantitative reverse transcription PCR (RT-qPCR) was used to measure the TgIF1 transcript levels in in the IF1^Ty^, IF1^KO^, and IF1^Over^ strains. Actin was utilized as a control housekeeping gene. Three technical replicates were used over five biological replicates for each strain. Expression levels were normalized to IF1^Ty^ using the 2^−ΔΔCt^ method. Unpaired, two-tailed t-test (p = 0.001 to 0.01: **, p < 0.0001: ****). **E.** Lysates from equivalent numbers of IF1^Ty^, IF1^KO^, and IF1^Over^ parasites were separated via SDS-PAGE then first probed with antibodies against Ty and HA. Membranes were later probed with antibodies against tubulin as a loading control. Data are representative of three biological replicates. **F.** Densitometric analysis of HA signal in IF1^Over^ parasites normalized to Ty signal in IF1^Ty^ parasites. Tubulin levels used as a loading control. Unpaired, two-tailed t-test (p = 0.01 to 0.05: *). **G.** IF1^Ty^, IF1^KO^, and IF1^Over^ parasites were transiently transfected with a plasmid encoding SOD2-GFP. Intracellular parasites were then fixed and stained for DAPI (blue) and either anti-Ty (IF1^Ty^ and IF1^KO^) or anti-HA antibodies (IF1^Over^) (red). Scale bar: 5µm.

Using RT-qPCR, we did not detect TgIF1 transcripts in the IF1^KO^ parasites, confirming the knockout of the gene in this strain. We also detected a significant increase in TgIF1 transcript levels in the IF1^Over^ strain compared to the IF1^Ty^ strain, confirming that addition of an exogenous copy of TgIF1 results in increased expression of the gene (Figure 1D). Subsequent Western blot analyses confirmed the presence of TgIF1 signal in the IF1^Ty^ strain at the appropriate molecular weight (13-14 kDa); and, as expected, TgIF1 signal was not detected in IF1^KO^ parasites (Figure 1E). An increase of TgIF1 signal was observed in IF1^Over^ lysates compared to IF1^Ty^ lysates, and densitometry analysis confirmed that TgIF1 protein levels were significantly increased by approximately five-fold in the IF1^Over^ strain as compared to the IF1^Ty^ strain (Figure 1, E and F).

Intriguingly, although a band at 13-14 kDa for TgIF1 was detected in the IF1^Over^ strain, the presence of an additional product with a molecular weight of ∼37 kDa was consistently observed as well (Figure 1E). While IF1 has been shown to form high molecular weight oligomers when chemically crosslinked (Cabezón *et al*., 2001; Faccenda *et al*., 2017; Gahura *et al*., 2018), to our knowledge this seems to be the first instance of a naturally stable oligomer resisting denaturation. To determine the protein composition of the ∼37 KDa band, we performed anti-Ty and anti-HA immunoprecipitations with the IF1^Ty^ and IF1^Over^ strains, respectively, then visualized the elution fractions via SDS-PAGE and silver staining. The indicated bands were then analyzed by mass spectrometry (Figure S1A). We identified several peptide hits associated with TgIF1 present in samples derived from IF1^Over^ parasites and absent in samples from IF1^Ty^ parasites (Figure S1B). Peptide hits associated with proteins exceeding 25 kDa (which corresponds to the approximate difference between the low and high molecular weight bands in the western blot from IF1^Over^ parasite lysates) were considered contaminants. Consequently, TgIF1 is the most probable candidate given its appropriately low molecular weight (Figure S1B). This finding suggests that the ∼37kDa product in the IF1^Over^ strain is a TgIF1 homo-oligomer.

We next sought to determine the localization of TgIF1 in each of our parasite strains. To do this, we transiently transfected them with a plasmid encoding SOD2-GFP, which targets GFP to the mitochondrial matrix (Pino *et al*., 2007), and performed immunofluorescence assays using anti-Ty and anti-HA antibodies. We observed the colocalization of TgIF1 with SOD2-GFP in IF1^Ty^ and IF1^Over^ parasites and, as expected, and we did not detect TgIF1 in IF1^KO^ parasites (Figure 1G). These observations confirm that TgIF1 localizes to the mitochondrion of the parasite, and that genetic manipulation of the protein does not impair its localization to this organelle. Together, our analyses validate the generation of the IF1^Ty^, IF1^KO^, and IF1^Over^ strains, suggest that TgIF1 overexpression results in a stable homo-oligomer, and confirm the mitochondrial localization of TgIF1.

### Overexpression of TgIF1 results in increased TgIF1 bound to the ATP synthase

We next sought to determine whether increased expression of TgIF1 impacts its interaction with the ATP synthase complex. To this end, we used two-dimensional blue native PAGE (2D BN-PAGE) to probe the subunit composition of the ATP synthase protein complex. Membranes containing IF1^Ty^ samples were incubated with anti-Ty antibodies (Figure 2A), while membranes containing IF1^Over^ samples were incubated with anti-HA antibodies (Figure 2B). Our results demonstrate that TgIF1 (13-14 kDa) interacts with the ATP synthase dimer (1860 kDa) in both the IF1^Ty^ and IF1^Over^ strains (Figure 2, A and B). Notably, the increased signal in the ATP synthase dimer area of the membrane in IF1^Over^ parasites compared to IF1^Ty^ suggests that TgIF1 overexpression increases TgIF1 binding to the ATP synthase (Figure 2, A and B).

**Figure 2.**
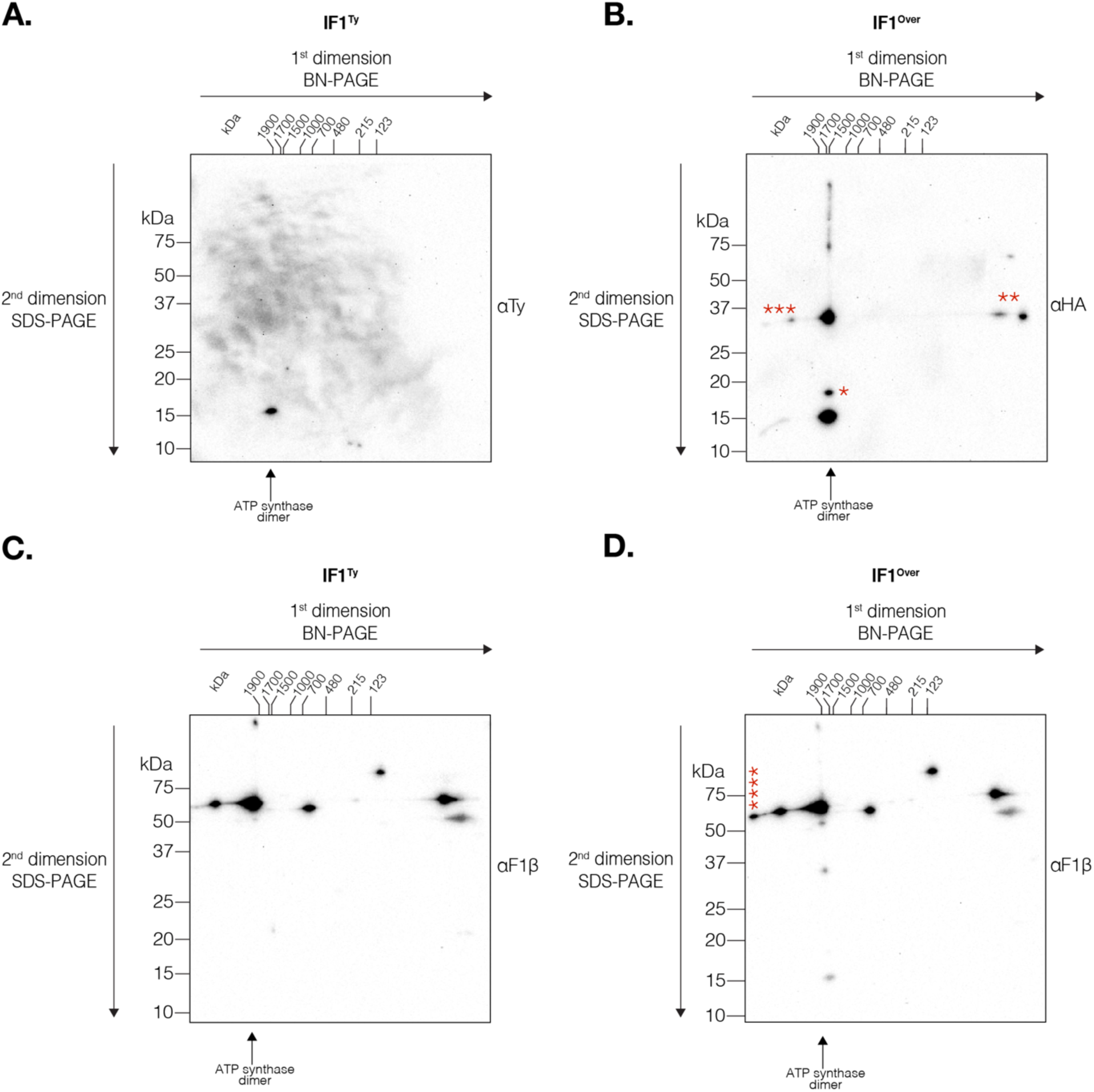
Overexpression of TgIF1 results in more IF1 bound to the ATP synthase. **A, B**. Lysates from (**A**) IF1^Ty^ and (**B**) IF1^Over^ parasites were resolved via two-dimensional blue native PAGE (2D BN-PAGE) then probed with antibodies against Ty (**A**) or HA (**B**) to assess the amount of TgIF1 bound to the ATP synthase in each strain. Membranes were exposed for the same amount of time. **C, D**. Both the IF1^Ty^ and IF1^Over^ membranes were stripped and re-probed with an antibody against the ATP synthase F1β subunit. Membranes were exposed for the same amount of time. Representative images of three biological replicates. Asterisks mark specific signals discussed in the main text.

Intriguingly, the high molecular weight TgIF1 oligomer (just below 37 kDa) previously observed by Western blot analysis (Figure 1E) is also associated with the ATP synthase dimer (Figure 2B). In addition, a signal (*) at a slightly higher molecular weight than the TgIF1 monomer signal (just below 20 kDa) was also detected on the IF1^Over^ immunoblot (Figure 2B). We have not previously observed a TgIF1 oligomer of this size via immunoblot analysis; its identity warrants further investigation. Lastly, although TgIF1 overexpression results in more TgIF1 bound to the ATP synthase dimer, not all of the overexpressed TgIF1 is able to bind, as two ∼37 kDa signals can be observed (**) at the low molecular weight end of the first-dimension ladder (Figure 2B).

Next, the anti-Ty and anti-HA antibodies were stripped from the IF1^Ty^ and IF1^Over^ membranes and each was re-probed with an antibody against the F1β subunit of the ATP synthase. (Figure 2, C and D). With the anti-F1β staining, several different assemblies of the *T. gondii* ATP synthase were detected on both immunoblots (Figure 2, C and D): high molecular weight oligomeric assemblies that inefficiently enter the gel (>1900 kDa), and potential ATP synthase dimers and monomeric assemblies (∼1860 and ∼700 kDa, respectively.) Additionally, we observed signals at ∼123 kDa and below. We speculate that the signal at ∼123 kDa is an hererodimer formed by the α and β subunits of the enzyme (Kane *et al*., 2010), and the low molecular weight signal (<123 kDa) represents β subunit monomers, which have a predicted molecular weight of 60 kDa. No TgIF1 signal was detected in molecular weight regions corresponding to the ATP synthase monomer or its assembly intermediates on the first-dimension gel (Figure 2, A-D), suggesting that TgIF1 exclusively binds to the dimeric form of the ATP synthase and not to its monomer or its assembly intermediates. Interestingly, 2D BN-PAGE analysis of IF1^Over^ lysates suggests that TgIF1 homo-oligomers can interact with larger ATP synthase oligomers (***) (Figure 2, B and D). Furthermore, an additional higher order ATP synthase assembly (****) was observed by F1β staining in two out of three replicates for IF1^Over^ parasites (Figure 2D), but not in any of the IF1^Ty^ replicates (Figure 2B). These data suggest that TgIF1 overexpression may increase the higher order oligomerization of the *T. gondii* ATP synthase.

Taken together, our 2D BN-PAGE experiments reveal that TgIF1 overexpression results in an increase amount of protein bound to the ATP synthase. Additionally, we show that both the monomeric and oligomeric forms of TgIF1 are capable of binding to the ATP synthase when TgIF1 is overexpressed. Lastly, we show that while TgIF1 is not able to bind to ATP synthase assemblies smaller than its dimeric form, TgIF1 homo-oligomer can bind to larger oligomers of the ATP synthase, and TgIF1 overexpression appears to promote the assembly of these larger oligomers.

### Disruption of TgIF1 results in the rewiring of gene regulatory networks in *T. gondii*

We next sought to understand the extent to which *T. gondii* parasites with altered levels of TgIF1 are different on a transcriptional level. To this end, we performed bulk RNA sequencing to generate transcriptomic data from each parasite line: 3 biological replicates for each (Figure 3A). We observed a strong correlation (R > 0.98) in gene expression between each biological replicate (Figure S2A) and principal component analysis revealed a separation between IF1^Ty^, IF1^KO^ and IF1^Over^ parasites based on their gene expression patterns (Figure 3B).

**Figure 3.**
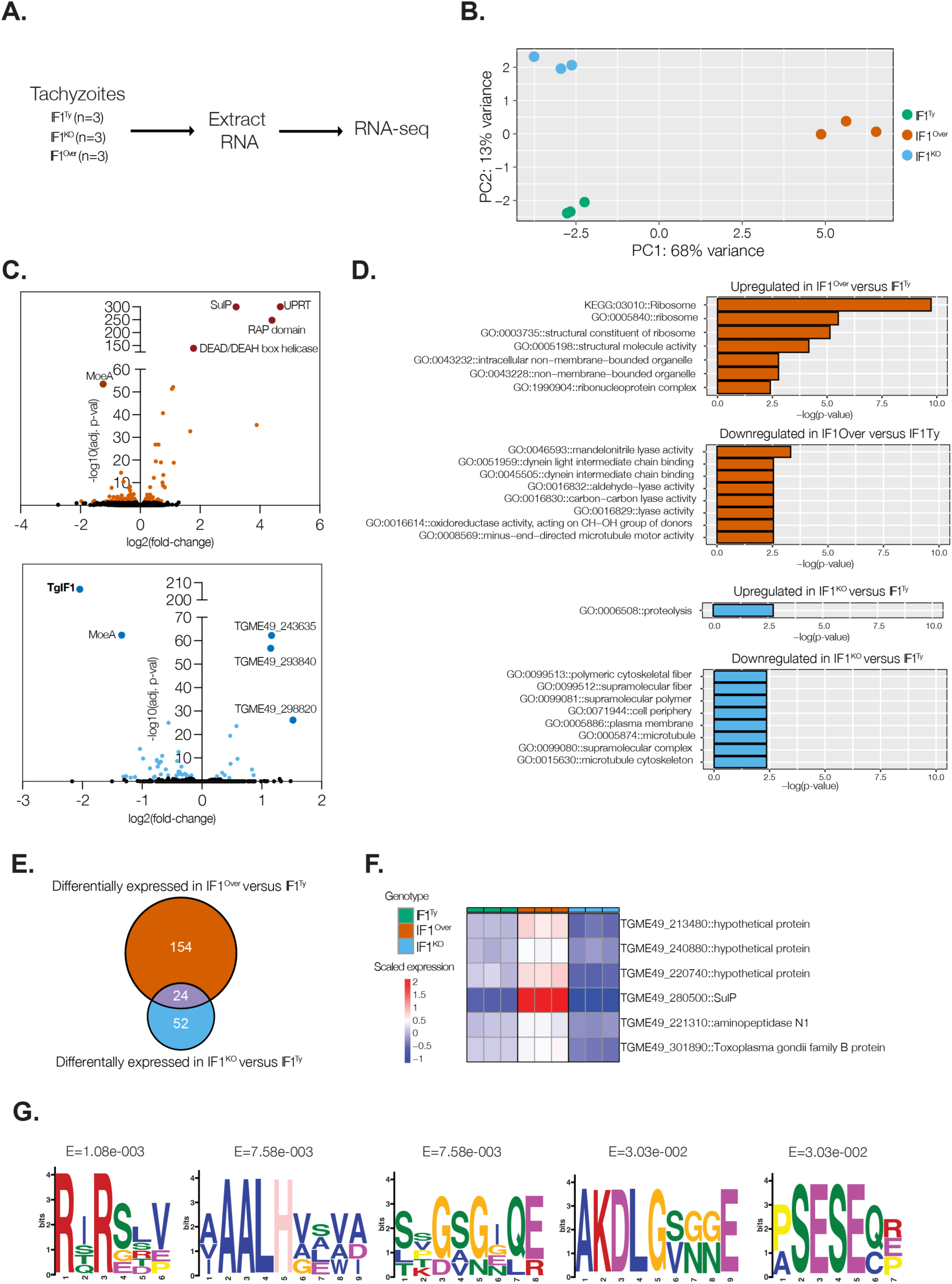
Transcriptomic analysis of parasites lacking and overexpressing TgIF1 reveals altered expression of genes associated with various biological processes. **A.** Schematic depicting the experimental design to generate the transcriptomic data from the three parasite lines. **B.** PCA plot displaying the variance explained in the first two principal components. Each data point represents one biological replicate. **C.** Volcano plots displaying the differentially expressed genes in the IF1^Over^ (upper) and IF1^KO^ strains (lower) compared to IF1^Ty^ parasites. Genes with a Benjamini and Hochberg adjusted p-value of < 0.05 were considered significantly different. **D.** Gene onthology (GO) Term enrichment analysis of genes with significantly different expression in IF1^Over^ strain (upper) and IF1^KO^ strain (lower) compared to IF1^Ty^ parasites. Bar plots represent gene sets containing 2 of more genes and a p value of < 0.01. **E.** Venn diagram displaying the unique and overlapping differentially expressed genes in the IF1^Over^ and IF1^KO^ parasites compared to IF1^Ty^ parasites. **F.** Heatmap displaying the scaled expression of genes that display increased expression in IF1^Over^ parasites and decreased expression IF1^KO^ compared to IF1^Ty^ parasites. **G.** Amino acid motifs enriched (Fisher’s exact test, E value < 0.05) in the genes that display increased expression in IF1^Over^ parasites and decreased expression IF1^KO^ compared to IF1^Ty^ parasites.

We compared the transcriptomes of IF1^KO^ and IF1^Over^ parasites to IF1^Ty^ parasites to assess the extent to which gene expression is altered due to the perturbation of TgIF1 levels. We identified 206 differentially expressed genes (adjusted p value < 0.05) when comparing IF1^KO^ and IF1^Over^ parasites to IF1^Ty^ parasites (Figure 3C). As expected, TgIF1 (TGME49_215350) was among the differentially expressed genes (Figure S2B). Gene Ontology (GO) term enrichment analysis revealed altered expression of genes associated with various cellular processes in IF1^KO^ and IF1^Over^ parasites compared to IF1^Ty^ parasites (Figure 3D; Table S1). We next sought to identify genes that displayed differential expression in both IF1^KO^ and IF1^Over^ parasites when compared to IF1^Ty^ parasites. We identified 24 genes differentially expressed in both in IF1^KO^ and IF1^Over^ parasites relative to IF1^Ty^ parasites (Figure 3E). Among these genes, six displayed gene expression changes that positively correlated with TgIF1 expression (Figure 3F). Their increased gene expression in IF1^Over^ parasites when compared to IF1^Ty^ parasites, and their decreased expression in IF1^KO^ parasites when compared to IF1^Ty^ parasites suggest that TgIF1 positively regulates these genes.

We also performed motif-based analysis on the set of genes positively correlated with TgIF1 expression (Figure 3F) to discover enriched motifs in the protein products of these genes. We identified six significantly enriched motifs (Fisher’s exact test, E value < 0.05) (Figure 3G, Table S2). The amino acid motif RIRSRV was significantly enriched (E =1.8×10^−3^) in each of the genes’ protein products (Figure 3G). These motifs might be critical for TgIF1-mediated regulation, via direct or indirect interaction, of this subset of genes. Together, our results reveal that both an increase and decrease in TgIF1 leads to distinct gene expression changes in parasites, and that TgIF1 may serve as a critical regulator of a subset of genes, in additional to its canonical role in the mitochondria.

### TgIF1 knockout results in decreased mitochondrial cristae density

In other systems, the role of IF1 in regulating cristae density remains controversial. While several groups have reported that IF1 overexpression increases cristae density, and its knock out reduces it (Campanella *et al*., 2008; Faccenda *et al*., 2017; Romero-Carramiñana *et al*., 2023), others investigations have not observed this (Fujikawa *et al*., 2012; Nakamura *et al*., 2013; Kahancová *et al*., 2020). Interestingly, one study even reported that IF1 overexpression decreased cristae density (Weissert *et al*., 2021). We therefore sought to investigate how the knockout of TgIF1 alters mitochondrial cristae density in *T. gondii*. To do so, we prepared samples from intracellular IF1^Ty^ and IF1^KO^ parasites for transmission electron microscopy. Quantification of mitochondrial cristae density revealed a statistically significant decrease in IF1^KO^ parasites compared to IF1^Ty^ (Figure 4, A and B). As there was no significant difference in the mitochondrial areas measured between the two strains (Figure 4C), these results indicate that TgIF1 plays a role in the maintenance of mitochondrial cristae density.

**Figure 4.**
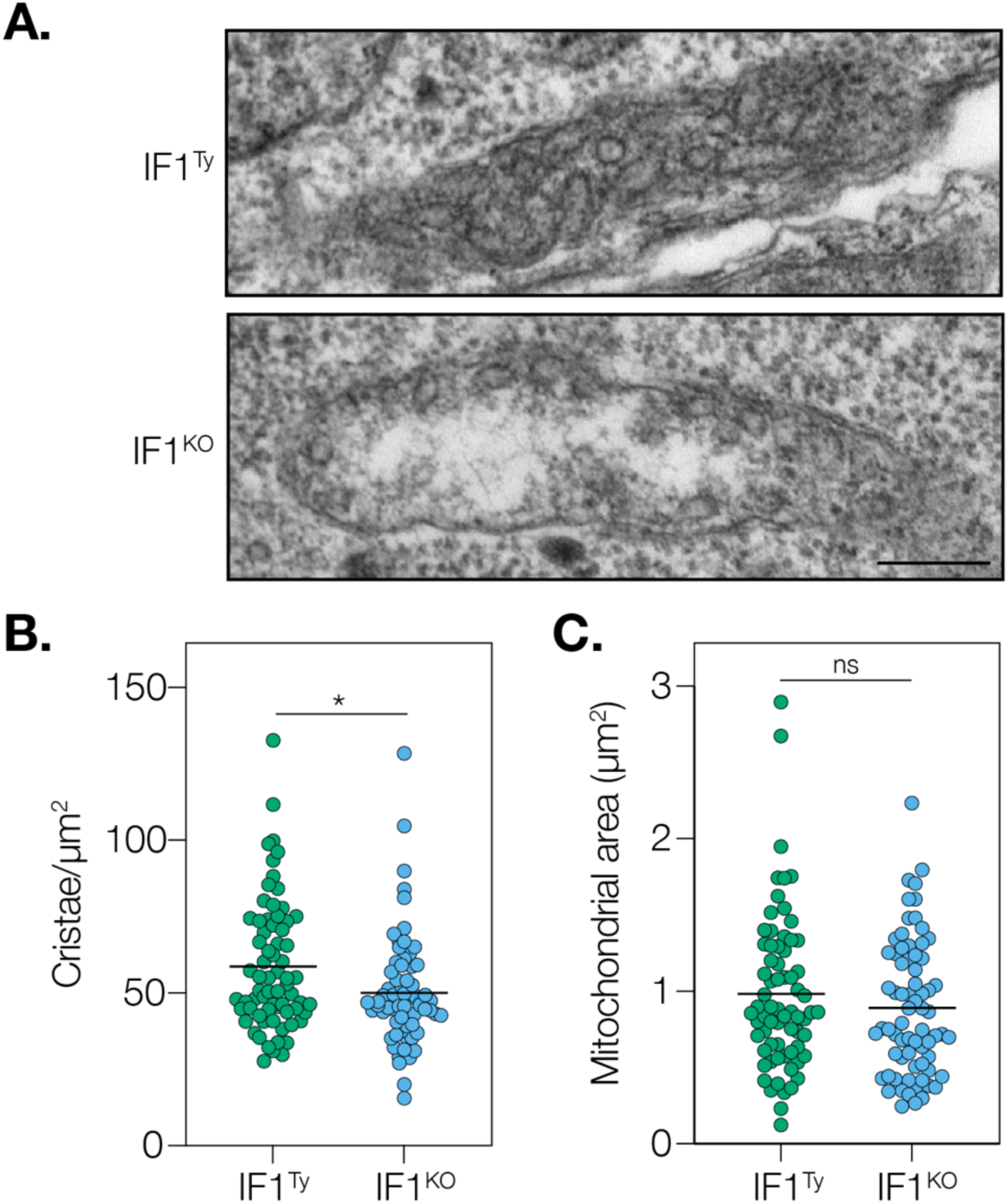
Mitochondrial cristae density is decreased in IF1^KO^ parasites. **A.** Representative electron micrographs of mitochondrial sections from IF1^Ty^ and IF1^KO^ parasites. Scale bar: 500nm. **B.** Quantification of cristae/µm^2^ of mitochondrial area from IF1^Ty^ and IF1^KO^ parasites. Data represent 70 sections of each strain. Unpaired, two-tailed t-test (p = 0.01 to 0.05: *). **C.** Mitochondrial areas (µm) measured from sections analyzed in (B). Unpaired, two-tailed t-test (ns = not significant).

### Effects of TgIF1 knockout and overexpression on ATP synthase dimerization, ATPase activity, metabolism, and membrane potential

We next investigated the extent to which altered TgIF1 expression affects ATP synthase dimerization. In yeast and mammals, how IF1 impacts ATP synthase dimerization is unclear. While several studies have shown a potential role of IF1 in ATP synthase dimerization (García *et al*., 2006; Campanella *et al*., 2008; Santacatterina *et al*., 2016; Romero-Carramiñana *et al*., 2023), other research indicates that IF1 does not influence dimerization of the enzyme complex (Tomasetig *et al*., 2002; Nakamura *et al*., 2013; Barbato *et al*., 2015). In *T. gondii,* structural studies have shown that TgIF1 homodimerizes via its C terminal region, allowing it to form an intra-dimeric bridge within each ATP synthase dimer of the larger hexameric ATP synthase oligomer observed in the parasite (Muhleip *et al*., 2021). This contrasts with what has been shown in mammalian cells, where IF1 forms inter-dimeric bridges within two separate ATP synthase dimers that make up a tetramer (Gu *et al*., 2019).

To investigate the role of TgIF1 in ATP synthase dimerization, we utilized BN-PAGE. Lysates were prepared from IF1^Ty^, IF1^KO^, and IF1^Over^ parasites, resolved by BN-PAGE, and processed for immunoblot analysis using antibodies against F1β. As there were no changes in the band representing the dimeric form of the ATP synthase between the strains (Figure 5A), we conclude that TgIF1 has no effect on ATP synthase dimerization in *T. gondii*.

**Figure 5.**
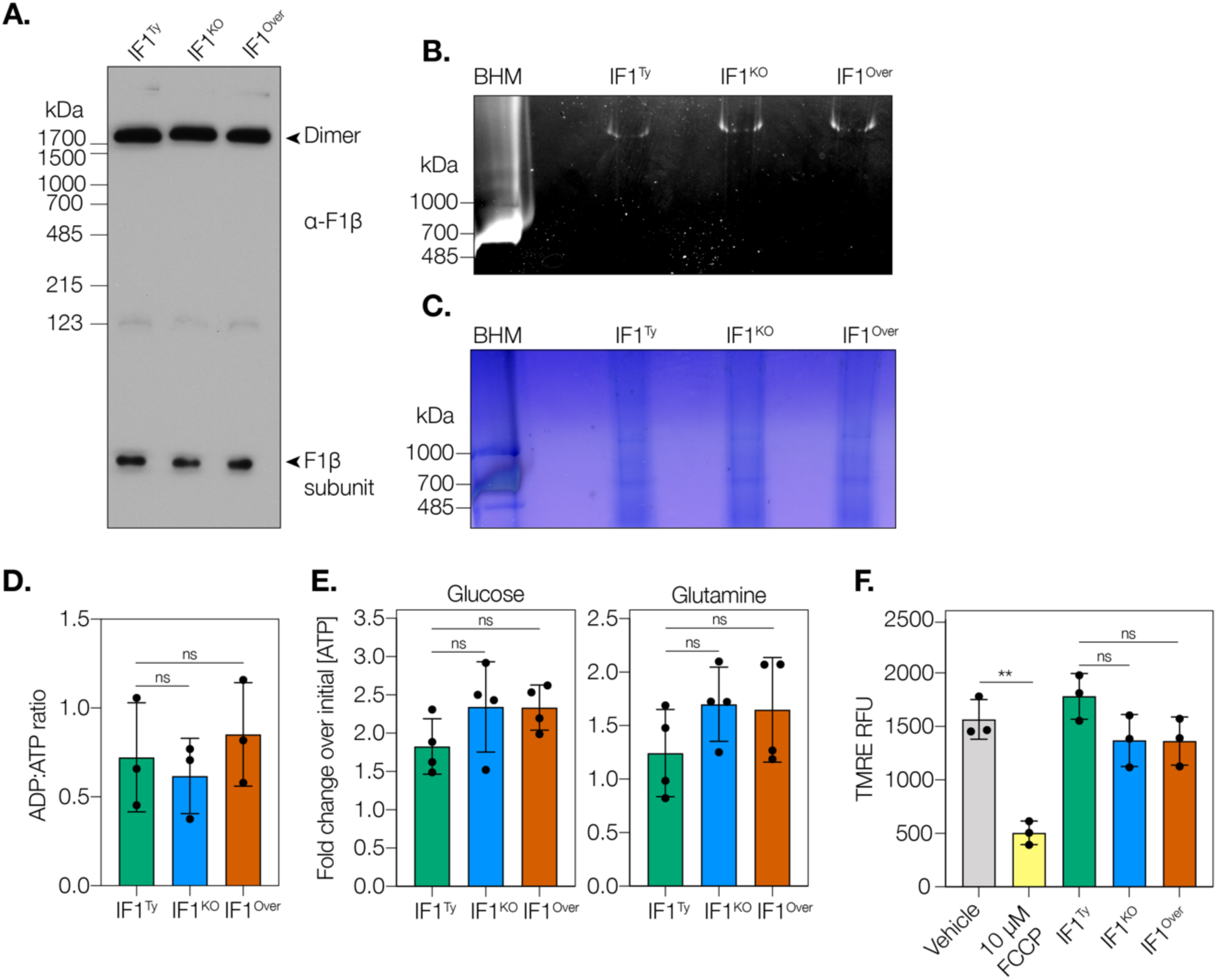
TgIF1 knockout and overexpression do not affect ATP synthase dimerization, ATPase activity, metabolism, or mitochondrial membrane potential under normal growth conditions. **A.** Lysates from IF1^Ty^, IF1^KO^, and IF1^Over^ parasites were resolved by blue native PAGE (BN-PAGE) and probed with an antibody against the ATP synthase F1β subunit. Data are representative of three biological replicates. **B.** Lysates from IF1^Ty^, IF1^KO^, and IF1^Over^ parasite strains were resolved by clear native PAGE (CN-PAGE) then subjected to in-gel ATPase activity assays. Bovine heart mitochondria (BHM) is used as a positive control. Data are representative of four biological replicates. **C.** Total Coomassie stain of gel shown in (**B**) to confirm loading of equivalent protein amounts. **D.** Cellular ADP:ATP ratios were determined from three biological replicates of IF1^Ty^, IF1^KO^, and IF1^Over^ parasites. Unpaired, two-tailed t-test (ns = not significant). **E, F**. Relative ATP concentrations of IF1^Ty^, IF1^KO^, and IF1^Over^ parasites. Parasites were incubated for 1 hour with 2-deoxy-D-glucose (2-DG) to inhibit glycolysis plus either (**E**) glucose or (**F**) glutamine. ATP levels for each condition were normalized to the initial ATP concentration of each strain. Data represent four biological replicates. Unpaired, two-tailed t-test (ns = not significant). **G.** Mitochondrial membrane potential measurements of the IF1^Ty^, IF1^KO^, and IF1^Over^ strains using TMRE. Data represent 3 biological replicates. Unpaired, two-tailed t-test (ns = not significant, p = 0.001 to 0.01: **).

Additionally, we wanted to investigate whether changes in TgIF1 protein levels affected the hydrolytic (ATPase) activity of the ATP synthase. To do this, we utilized a previously developed in-gel ATPase assay that detects ATPase activity through the formation of white lead phosphate precipitates (Suhai *et al*., 2009). We prepared clear native PAGE samples from IF1^Ty^, IF1^KO^, and IF1^Over^ parasites as well as from bovine heart mitochondria (BHM), which served as both a positive control and a molecular weight marker for the assay. The BHM sample formed a white precipitate around 700 kDa (Figure 5B). Samples from IF1^Ty^, IF1^KO^ and IF1^Over^ parasites resulted in weaker bands at a much higher molecular weight (Figure 5B). Equivalent protein loading between the parasite samples was confirmed through Coomassie staining of the gel (Figure 5C). These results suggest that the knockout and overexpression of TgIF1 do not have significant effects on *T. gondii* ATPase activity.

We were also interested in investigating broader metabolic changes in the IF1^Ty^, IF1^KO^, and IF1^Over^ parasite strains. We first investigated the ADP:ATP ratio, which has been shown to be an important indicator of energy status within the cell (Yuan *et al*., 2013). We did not observe any significant changes in ADP:ATP ratios when TgIF1 was knocked out or overexpressed (Figure 5D), suggesting that TgIF1 does not alter this aspect of parasite energy metabolism. To more specifically investigate changes to ATP originating from various metabolic sources, we next utilized a previously described assay (Huet *et al*., 2018). Briefly, parasites were incubated with a glycolysis inhibitor and provided with either sufficient glucose to overcome glycolytic inhibition, or sufficient glutamine to promote oxidative phosphorylation (OXPHOS). While glucose contributes to ATP production via both glycolysis and OXPHOS, glutamine is only able to contribute to ATP production via OXPHOS (MacRae *et al*., 2012). Using this assay, we found that the amount of ATP produced from glucose or glutamine in IF1^KO^ and IF1^Over^ parasites was similar to the amount produced in the IF1^Ty^ strain (Figure 5E). These data suggest that the modulation of TgIF1 levels through knockout and overexpression has no significant effects on ATP production in *T. gondii*.

Aside from its role in energy production, the ATP synthase also plays an important role in mitochondrial membrane potential maintenance. In other systems, IF1 has been shown to impact mitochondrial membrane potential through its binding and inhibition of the ATP synthase. More specifically, increases in IF1 protein are correlated with membrane potential hyperpolarization (Sanchez-Cenizo *et al*., 2010; Esparza-Moltó *et al*., 2021). It has also been shown that IF1 ablation can cause membrane potential depolarization (Esparza-Moltó *et al*., 2021). We thus decided to investigate whether TgIF1 knockout or overexpression had any effect on mitochondrial membrane potential in *T. gondii*. Syringe-released parasites were stained with the lipophilic cationic dye TMRE to assess mitochondrial membrane potential via fluorescent readout. FCCP, a protonophore that dissipates membrane potential, was included in the assay as an positive control. Using this method, we did not find any significant differences in membrane potential between IF1^Ty^, IF1^KO^, and IF1^Over^ parasite strains (Figure 5F). Together, these data suggest that TgIF1 does not have significant effects on *T. gondii* ATP synthase dimerization, ATPase activity, metabolic flux, or mitochondrial membrane potential.

### TgIF1 levels are critical for responding to hypoxia

In metazoan systems, the pH-based control of IF1 activity allows for ATP synthase inhibition to be regulated by mitochondrial stressors, such as hypoxia, which result in matrix acidification (Gore *et al*., 2022). To investigate a potential role of TgIF1 in hypoxia, we compared parasite growth under normoxic (21% O_2_) and hypoxic (0.5% O_2_) conditions. Parasites from IF1^Ty^, IF1^KO^, and IF1^Over^ strains were grown undisturbed on HFF monolayers under hypoxic or normoxic conditions for eight days before fixation and staining with crystal violet. Hypoxic conditions reduced plaque size and number for all three strains, but the reduction was greater in the IF1^KO^ and IF1^Over^ parasites (Figure 6A). We quantified both the number of plaques and the size of plaques for each strain under both conditions, then normalized the values at 0.5% O_2_ to the values at 21% O_2_ for each strain to illustrate strain-specific differences. Our results show that the decreases in plaque numbers due to hypoxia or strain-specific differences were not statistically significant (Figure 6, B and C). However, when plaque size was quantified, the decreases under hypoxic conditions compared to normoxic conditions were statistically significant in all three strains (Figure 6D). Further, when the size of plaques in 0.5% O_2_ was normalized to the size at 21% O_2_, the IF1^KO^ and IF1^Over^ strains had significantly smaller sizes in comparison to the IF1^Ty^ strain (Figure 6E). Together, our data show that perturbations to TgIF1 levels, whether through knockout or overexpression, negatively impact the ability of the parasites to grow under hypoxic conditions.

**Figure 6.**
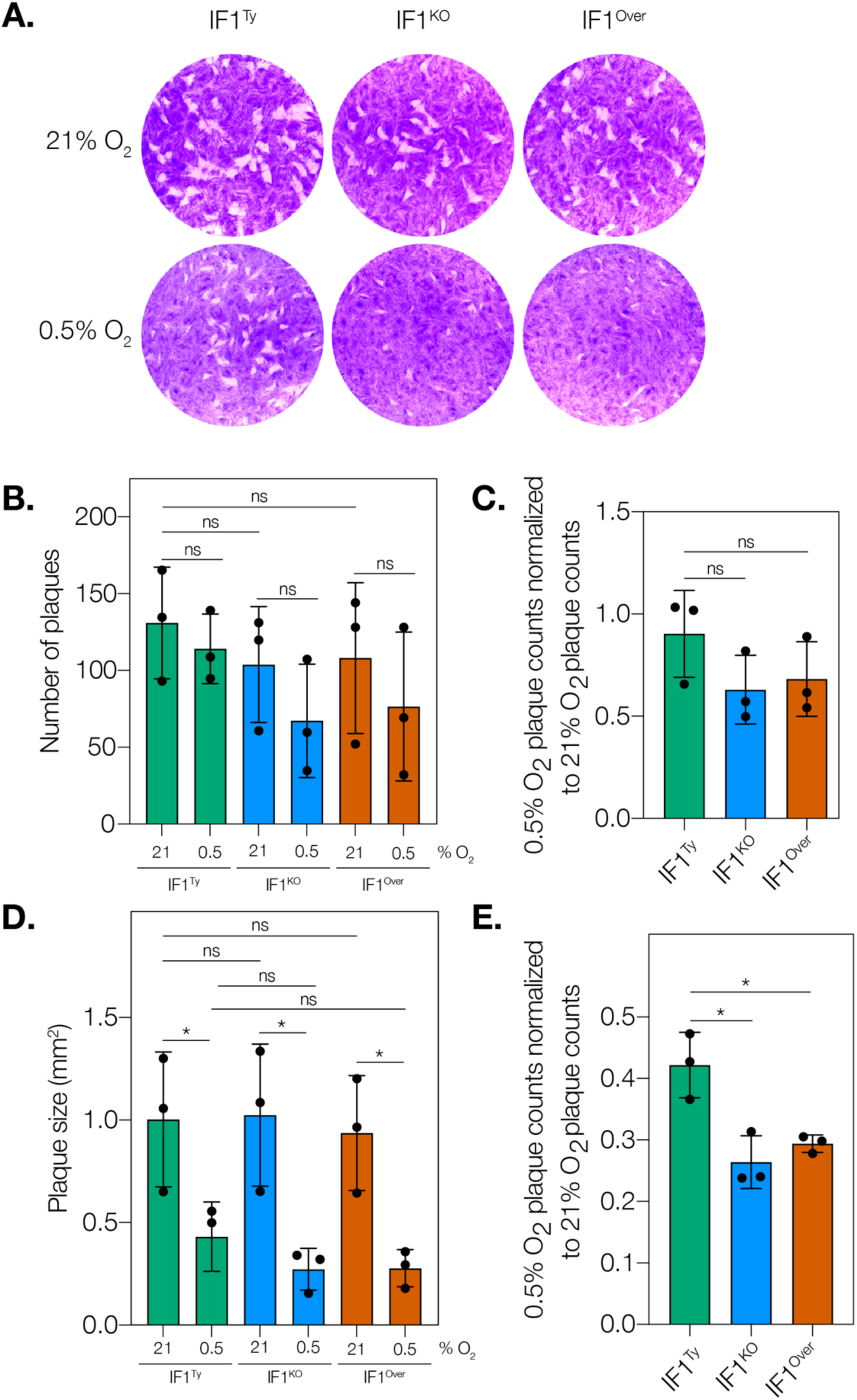
Knockout and overexpression of TgIF1 decrease parasite replication under hypoxic conditions. **A.** Plaque assay of the IF1^Ty^, IF1^KO^, and IF1^Over^ strains grown under normoxic (21% O_2_) or hypoxic (0.5% O_2_) conditions. Data are representative of three biological replicates. **B.** Quantification of the plaque numbers per well in (**A**). Paired two-tailed t-tests were conducted to compare the same strain under different conditions, and unpaired t-tests were used for comparisons between different strains. (ns = not significant). **C.** Average plaque numbers at 0.5% O_2_ from (**B**) were normalized to plaque numbers at 21% O_2_ for each parasite strain. Unpaired, two-tailed t-test (ns = not significant). **D.** Plaque size was manually measured for 20 plaques in each well. Paired two-tailed t-tests were conducted to compare the same strain under different conditions, and unpaired t-tests were used for comparisons between different strains. (ns = not significant, p = 0.01 to 0.05: *). **E.** Average plaque size at 0.5% O_2_ from (D) was normalized to plaque size at 21% O_2_ for each parasite strain. Upaired, two-tailed t-test (p = 0.01 to 0.05: *).

### A potential role of TgIF1 in the mitohormetic response of *T. gondii*

Given the established role of IF1 in mitohormesis (Yun and Finkel, 2014), we investigated whether TgIF1 could trigger a similar response in *T. gondii*. To evaluate this, we utilized monensin: an Na^+^/H^+^ exchanging ionophore that has been shown to damage the *T. gondii* mitochondrion through induction of ROS release (Charvat and Arrizabalaga, 2016). We reasoned that increased levels of TgIF1 in the IF1^Over^ strain could potentially induce a mitohormetic response in *T. gondii*, and wanted to see whether IF1^Ty^, IF1^KO^, and IF1^Over^ parasites respond differently to the ROS stress associated with monensin treatment. To determine strain-specific differences, we performed a plaque assay with each strain in the presence of monensin or a vehicle control for 24 hours before removal and replacement with fresh, conventional culture media. Monensin treatment caused significant decreases in the number of plaques compared to vehicle control for all three strains (Figure 7, A and B). However, when the number of plaques from monensin-treated wells was normalized to vehicle, there were no significant differences between the three strains (Figure 7C). When plaque size was measured, monensin caused a significant decrease in all three strains compared to vehicle, but this decrease was of a lower magnitude in the IF1^Over^ strain (Figure 7D). Further, when the size of plaques in monensin-treated wells was normalized to that of vehicle-treated wells, IF1^Over^ parasites showed a significant increase in plaque size in comparison to IF1^Ty^ and IF1^KO^ strains (Figure 7E). Taken together, these data suggest that increased TgIF1 levels might promote a mitohormetic response in *T. gondii*, allowing the parasite to mitigate the ROS stress induced by monensin treatment.

**Figure 7.**
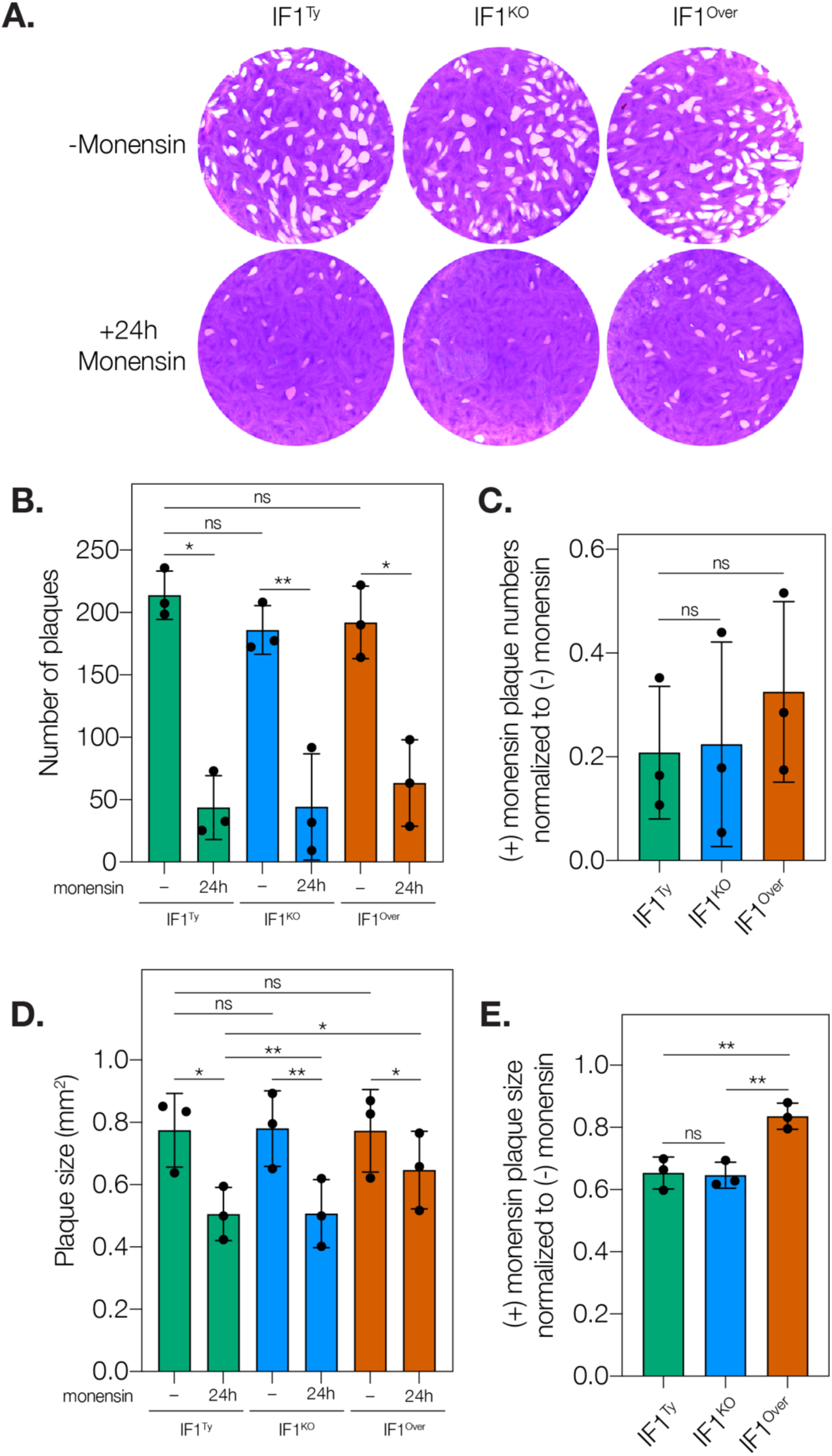
Overexpression of TgIF1 increases parasite growth following incubation with monensin. **A.** 500 parasites from IF1Ty, IF1KO, and IF1 Over strains were allowed to invade an HFF monolayer for two hours prior to a 24-hour treatment with 0.003 µM monensin or vehicle, 70% ethanol (−). After washout, parasites were allowed to grow undisturbed for a total of 7 days in normal growth medium. Data are representative of three biological replicates. **B.** Quantification of the plaque numbers per well. Paired two-tailed t-tests were conducted to compare the same strain under different conditions, and unpaired t-tests were used for comparisons between different strains. (ns = not significant, p = 0.001 to 0.01: *, p = 0.0001 to 0.001: **). **C.** Average plaque numbers from monensin-treated samples from (**B**) were normalized to plaque numbers from non-treated samples for each parasite strain. Paired two-tailed t-tests were conducted to compare the same strain under different conditions, and unpaired t-tests were used for comparisons between different strains (ns = not significant). **D.** Plaque size was manually measured for 20 plaques in each well using Fiji. Paired two-tailed t-tests were conducted to compare the same strain under different conditions, and unpaired t-tests were used for comparisons between different strains. (ns = not significant, p = 0.001 to 0.01: *, p = 0.0001 to 0.001: **). **E.** Average plaque size from monensin-treated samples (D) was normalized to plaque size from non-treated samples for each parasite strain. Unpaired, two-tailed t-test (ns = not significant, p = 0.001 to 0.01: **).

## DISCUSSION

Recent work has uncovered that the apicomplexan mitochondrion contains a high concentration of essential, phylum-specific proteins, thus making it an ideal target for the development of novel therapeutics (Usey and Huet, 2022; Lamb *et al*., 2023). The energy-generating mitochondrial ATP synthase is particularly divergent in the model apicomplexan *T. gondii*: over half of its subunits have no known homologs outside of the phylum (Huet *et al*., 2018; Salunke *et al*., 2018; Maclean *et al*., 2021; Muhleip *et al*., 2021). While we are beginning to uncover the roles of some of these divergent subunits (Muhleip *et al*., 2021; Usey and Huet, 2023), the regulatory mechanisms governing this critical complex are largely unexplored. In the present study, we characterized the *T. gondii* homolog of IF1, a conserved ATP synthase inhibitor.

Intriguingly, when we overexpressed TgIF1, we observed the formation of a stable, high molecular weight oligomer that exhibited remarkable stability. While the TgIF1 monomer is approximately 13-14 kDa, the high molecular weight oligomer was approximately 37 kDa. Immunoprecipitation followed by silver staining and mass spectrometry suggests that this high molecular weight oligomer is a TgIF1 homo-oligomer. Studies in other systems have observed that IF1 will form high molecular weight homo-oligomers when chemically crosslinked (Cabezón *et al*., 2001; Faccenda *et al*., 2017; Gahura *et al*., 2018). Such crosslinking studies have revealed that the IF1 dimer runs between 20-25 kDa, the trimer at ∼37 kDa, and the tetramer at ∼50 kDa in mammals (Faccenda *et al*., 2017; Gahura *et al*., 2018). Interpreting our results based on these data, it appears that the stable, high molecular weight (∼37 kDa) oligomer we observe in the IF1^Over^ line may be a TgIF1 trimer. However, the reasons behind its apparent ability to resist denaturation prior to SDS-PAGE remain unknown.

In other systems, the form of IF1 that binds to ATP synthase is either a monomer or dimer, depending on the organism (Cabezón *et al*., 2001; Gledhill *et al*., 2007; Robinson *et al*., 2013; Le Breton *et al*., 2016; Gahura *et al*., 2018). Cryo-electron microscopy studies of the *T. gondii* ATP synthase showed a TgIF1 dimer bound to the ATP synthase dimer (Muhleip *et al*., 2021). These results suggest that the inhibitory form of IF1 in *T. gondii* is the dimer, as is the case in metazoans and in the parasitic kinetoplastid *Trypanosoma brucei* (Cabezón *et al*., 2001; Panicucci *et al*., 2017; Gahura *et al*., 2018). Thus, whether the putative TgIF1 trimer we observe binding to the ATP synthase in our 2D BN-PAGE experiments is physiologically relevant outside of our exogenous overexpression system remains to be determined. Nevertheless, our results suggesting that an IF1 trimer can associate to the ATP synthase are, to our knowledge, the first of their kind.

When looking at transcriptional changes caused by the absence or the overexpression of TgIF1, we observe that perturbation of this protein leads to transcriptome-wide changes in *T. gondii*. Gene Ontology term analysis show an altered expression of genes linked to ribosomal activity, metabolism, and cytoskeletal function in IF1^KO^ and IF1^Over^ parasites compared to IF1^Ty^ parasites. Although the relationship between cytoskeletal function and TgIF1 expression has not been previously described, our analysis suggest a role for TgIF1 in transcriptional regulation and metabolism, reminiscent of the established link between IF1 and mitohormesis in other organisms (Yun and Finkel, 2014; García-Aguilar and Cuezva, 2018). Additionally, the IF1^Over^ dataset showed a significant enrichment of genes related to gene expression regulation, supporting the notion that TgIF1 overexpression can re-program gene expression and potentially contributes to a mitohormetic response in *T. gondii*.

The transcriptomic data also provided additional insights into the metabolic changes resulting from TgIF1 knockout and overexpression. For example, a MoeA domain-containing protein was significantly downregulated in both IF1^KO^ and IF1^Over^ parasite strains compared to IF1^Ty^. Proteins containing MoeA domains are important cofactors for enzymes involved in metabolism of sulfur, carbon, and nitrogen (Mendel and Bittner, 2006). Additional experiments using metabolomics may provide more detail on any metabolic effects associated with TgIF1. Other interesting candidates include a sulfate permease (SulP) family protein, a DEAD/DEAH box helicase, and a RAP domain-containing protein. SulP proteins are involved in the transport of various ions, including chloride, sulfate, hydroxyl, bicarbonate, and oxalate across cellular membranes (Mount and Romero, 2004). Additionally, the DEAD/DEAH box helicase has functions beyond its RNA unwinding activity; genes in this family have been shown to act in transcriptional regulation (Fuller-Pace, 2006). Lastly, RAP (RNA-binding domain abundant in apicomplexans) domain-containing proteins have been shown to interact with RNA to mediate a range of cellular functions (Lee and Hong, 2004). Characterization of these candidates might yield further insight into the roles of TgIF1. The RNA-sequencing dataset generated in this study has also opened new avenues of inquiry linked to the regulatory role of TgIF1. Two of the top five differentially expressed genes in the IF1^KO^ dataset encode hypothetical proteins that had no structural homologs as determined by HHPRED (Zimmermann *et al*., 2018). Future investigations into the localization and function of these gene products could reveal important information on the role of TgIF1 within the parasites. Similarly, motif analysis of genes upregulated by TgIF1 identified six potential regulatory motifs. These motifs imply a potential direct or indirect regulatory role for TgIF1. To confirm our hypothesis, functional studies, including mutagenesis of these motifs, will be required.

In addition, we observed that TgIF1 overexpression increased the higher order oligomerization of ATP synthase complexes, while both TgIF1 knockout and overexpression left the dimeric form unaffected. Despite structural evidence suggesting that TgIF1 contributes to the dimerization of the *T. gondii* ATP synthase by forming an intradimeric bridge (Muhleip *et al*., 2021), the lack of a dimerization phenotype upon TgIF1 loss or overexpression might be attributable to the presence of an extensive dimerization interface in this enzyme. This dimerization interface has been shown to be considerably larger than the interface in mammals and yeast. More specifically, the *T. gondii* dimerization interface is composed of eleven different subunits from each monomer, many of which are apicomplexan-specific, that extend deep into the structure (Muhleip *et al*., 2021). As a result, the loss or overexpression of one dimerization component may not have any effect on the extremely stable interface. Nonetheless, disruptions to higher order oligomeric forms of the *T. gondii* ATP synthase have been observed following deletion of an ATP synthase subunit, despite a detectable impact on dimerization. Notably, these disruptions were associated with defects in mitochondrial cristae formation (Muhleip *et al*., 2021). Indeed, we observed a significant decrease in mitochondrial cristae density in the IF1^KO^ strain compared to the IF1^Ty^ strain. Taken together, our data suggest that TgIF1 plays a critical role in higher order ATP synthase oligomerization, with downstream impacts on cristae formation.

The role of IF1 in both ATP synthase oligomerization and cristae formation has been hotly contested in the literature (García *et al*., 2006; Campanella *et al*., 2008; Fujikawa *et al*., 2012; Nakamura *et al*., 2013; Barbato *et al*., 2015; Santacatterina *et al*., 2016; Faccenda *et al*., 2017; Kahancová *et al*., 2020; Weissert *et al*., 2021; Domínguez-Zorita *et al*., 2023). However, recent work may shed light on these apparent differences. A 2023 study suggests that there are two separate forms of ATP synthase in a cell: an active form and an inactive, or “sluggish”, IF1-bound form (Romero-Carramiñana *et al*., 2023). The authors of the study suggest that IF1 increases ATP synthase oligomerization and the “sluggish”, oligomeric assemblies of ATP synthase cluster at cristae tips, helping to shape cristae and create microdomains of membrane potential hyperpolarization. They also hypothesize that the “sluggish” ATP synthase represents a reservoir of enzyme that can be activated to meet higher energy needs upon demand and serves as a generator of ROS for cellular signaling (Romero-Carramiñana *et al*., 2023).

We also undertook a variety of biochemical and metabolic experiments to characterize the function of TgIF1. To first investigate whether TgIF1 had any effect on the hydrolytic activity of the ATP synthase, we conducted in-gel ATPase assays. We were unable to detect any difference in ATPase activity when TgIF1 was knocked out or overexpressed. We thus utilized ADP:ATP ratio assays, as well as glucose/glutamine ATP production assays, to see if we could observe any effects on other aspects of metabolism. We did not observe any changes in metabolism in our parasite strains. Similarly, mitochondrial membrane potential did not seem to change when TgIF1 was knocked out or overexpressed.

Although many studies have shown that increased IF1 levels result in decreased OXPHOS and mitochondrial membrane potential hyperpolarization due to blockage of proton backflow through the ATP synthase (Sanchez-Cenizo *et al*., 2010; Formentini *et al*., 2012; García-Aguilar and Cuezva, 2018; Esparza-Moltó *et al*., 2021), this prevailing school of thought has been challenged by several studies. Specifically, some groups have observed the opposite: mitochondrial membrane potential depolarization and increased OXPHOS occurred when IF1 is overexpressed (Weissert *et al*., 2021), and similarly, membrane potential hyperpolarization and decreased OXPHOS occurred when IF1 was ablated (Fujikawa *et al*., 2012; Barbato *et al*., 2015; Zhong *et al*., 2022). Further, others have found IF1 ablation to have no effect on membrane potential and OXPHOS (Galber *et al*., 2023). Potential explanations for such disparate observations may lie in the different cell types used, which differ in intrinsic IF1 content, and thus react differently to perturbations of IF1 levels (Romero-Carramiñana *et al*., 2023). Furthermore, explanations for the different observations may be due to whether they were gained from transient versus stable IF1 expression systems (Fujikawa *et al*., 2012; Barbato *et al*., 2015). As such, many of the reported observations may be due to the cells adapting to changes in IF1 levels, and may not reflect the behavior of cells that have already adapted to altered IF1 levels (Fujikawa *et al*., 2012). In the case of *T. gondii,* the observed lack of changes to metabolism and membrane potential may be due to the parasite’s inherent metabolic flexibility (MacRae *et al*., 2012), and an adaptation to the stable TgIF1 knockout and overexpression systems used in our study. Consequently, the generation of conditional TgIF1 knockdown and overexpression strains could provide insight into potential adaptations to TgIF1-mediated metabolic changes.

Aside from its proposed role in metabolism and mitochondrial cristae maintenance, IF1 has also been suggested to be critical for regulating the cellular response to hypoxia (Gore *et al*., 2022). Interestingly, we found that TgIF1 plays a role in the parasite’s ability to replicate under low oxygen conditions. We found that both TgIF1 knockout and overexpression decreased the growth of *T. gondii* tachyzoites under hypoxic conditions but had no effect on growth under normoxic conditions. In other systems, IF1 is important for the balance between preserving cellular energy and preserving the mitochondrial membrane potential. Under low oxygen conditions, the lack of a final electron acceptor for the electron transport chain can lead to mitochondrial membrane potential depolarization and acidification of the matrix (Gore *et al*., 2022). When this occurs, the ATP synthase can act in reverse, hydrolyzing ATP to pump protons back into the intermembrane space and preserve the mitochondrial membrane potential, the loss of which can result in cellular death (Gore *et al*., 2022). In the case of TgIF1, our data suggest that this balance cannot be perturbed: loss of TgIF1 (IF1^KO^) might result in futile and excessive wastage of ATP due to uninhibited reverse cycling of the ATP synthase, and that too much TgIF1 (IF1^Over^) prevents the maintenance of membrane potential, leading to detrimental downstream effects on mitochondrial health in hypoxic conditions.

Though low oxygen is one form of stress that IF1 has been shown to mediate, it has also been shown to regulate responses to a wide range of stressors through the process of mitohormesis (García-Aguilar and Cuezva, 2018). Our findings in *T. gondii* also support this notion, as parasites overexpressing TgIF1 were better able to divide and replicate following treatment with monensin. In mammals, the addition of a stressor was necessary to tease out the effect of IF1 overexpression (Formentini *et al*., 2014, 2017; Santacatterina *et al*., 2016). Providing support for this observation, a previously conducted CRISPR screen in *T. gondii* for genes involved in oxidative stress responses found that parasites with a disrupted TgIF1 locus were negatively impacted in their ability to respond to hydrogen peroxide stress. Thus, TgIF1 was given a negative phenotype score (−0.8) (Chen *et al*., 2021). Under baseline growth conditions, TgIF1 has a positive phenotype score (+1.8), indicating that the gene is not essential for survival under normal conditions (Sidik *et al*., 2016). These data, in combination with our own work, suggest that the application of stressors, such as ROS or hypoxic stress, will help to further unveil the role of TgIF1 in mitohormesis.

While our work has shed light on the phenotype associated with manipulations of TgIF1 levels, several important questions remain. One such question is whether TgIF1 can oligomerize into other forms and whether the putative TgIF1 trimer we observed is physiologically relevant. Other remaining questions include how the binding of TgIF1 to the ATP synthase is regulated, whether it plays a role in responding to other types of stressors, and whether it plays an important role in the slow-replicating bradyzoite form of *T. gondii*.

In summary, our work characterizing TgIF1 demonstrates that it is dispensable for ATP production and parasite survival under baseline growth conditions. Our work also demonstrates that TgIF1 plays a role in cristae morphology, the parasite’s response to hypoxia and other stressors, as well as gene expression regulation. The precise mechanism by which TgIF1 shields parasites from oxidative stress requires further investigation. Previous research has linked impaired ROS stress responses to reduced *T. gondii* virulence in knockout mice (Chen et al., 2021). Although a recent CRISPR-based screen in mice suggest that TgIF1 does not affect parasite virulence (Giuliano *et al*., 2024), it is crucial to determine whether TgIF1 is essential for parasite survival and development *in vivo*. Given the dramatically different environments *T. gondii* encounters throughout its complex life cycle, understanding how TgIF1 regulates ATP synthase activity will not only provide insights into this evolutionarily conserved process, it could also potentially identify novel therapeutic targets against apicomplexan parasites.

## MATERIALS AND METHODS

### Parasite culture

RH/TATi/Δku80 (Sheiner *et al*., 2011) tachyzoites and their derivatives were maintained in human foreskin fibroblasts (HFFs) (ATCC, cat. no. SCRC-1041). Strains were cultured at 37°C and 5% CO_2_ in DMEM supplemented with 2mM glutamine (GeminiBio, cat. no. 400-106) and 3% heat-inactivated fetal calf serum (IFS).

### Cloning and parasite strain generation

To generate the IF1^Ty^ strain, the pU6-Universal plasmid (Addgene, cat. no. 52694) was digested using the BsaI restriction enzyme then an sgRNA targeting the C terminus of TgIF1 (TGME49_215350) (P1 and P2) was inserted into the plasmid via Gibson assembly. A forward and reverse repair template, encoding a single Ty epitope tag with homology to the C terminus and 3’ UTR of TgIF1, was duplexed and dialyzed to reduce salt content (P3 and P4). 30µg of this repair template and 100µg of the pU6-Universal plasmid encoding Cas9 and the sgRNA were transfected into RH/TATi/Δku80 parasites as previously described (Sidik *et al*., 2014). After recovery from transfection, the population was subcloned via serial dilution and clonal lines were screened for correct integration of the Ty tag via PCR using P5 and P6. Expression of the Ty tag was confirmed via immunofluorescence and western blotting.

To create a complete knockout of the TgIF1 gene (IF1^KO^), sgRNAs targeting the N and C termini of the Ty-tagged gene (P7 and P8 for the N terminus; P9 and P10 for the C terminus) were each inserted into BsaI-digested pU6-Universal plasmids using Gibson assembly. The repair template encoding a pyrimethamine-resistant copy of the DHFR cassette was amplified from the DHFR-SAG4-TetO7-3xTy plasmid (a generous gift from Silvia Moreno) using P11 and P12, which have homology to the 5’ and 3’ UTRs of TgIF1, respectively. Approximately 15µg of this repair template and 50µg of each of the two pU6-Universal plasmids encoding Cas9 and the N and C terminal sgRNAs were transfected into IF1^Ty^ parasites as previously described (Sidik *et al*., 2014).

Pyrimethamine (Sigma Aldrich, cat. no. 46706-250MG) was used at 3µM to select for parasites containing the correct IF1^KO^ integration. After recovery from drug selection, the population was subcloned via serial dilution and clonal lines were screened for replacement of the TgIF1 locus with the DHFR cassette via PCR using P5 and P13. Knockout of the Ty-tagged TgIF1 was confirmed via immunofluorescence and western blotting.

For creation of the IF1 overexpression strain (IF1^Over^), a plasmid containing the Tub8 promoter, the TgIF1 CDS, a C-terminal in-frame single HA epitope tag, and the 3’ UTR of TGME49_231410 (ATP synthase b subunit) was assembled via Gibson assembly. From this plasmid, the repair template encoding Tub8-TgIF1-HA-ICAP2 3’ UTR was amplified using P14 and P15, which have homology to the *T. gondii* uracil phosphoribosyltransferase (UPRT) locus (TGME49_312480). An sgRNA targeting the UPRT locus (P16 and P17) was inserted into the BsaI-digested pU6-Universal plasmid using Gibson assembly. Approximately 25µg of this repair template and 50µg of the pU6-Universal plasmid encoding Cas9 and the UPRT sgRNA were transfected into IF1^KO^ parasites as previously described (Sidik *et al*., 2014). The thymidylate synthase inhibitor 5-fluoro-2’-deoxyuridine (FUDR) (Sigma Aldrich, cat. no. F0503-100MG) was used at 5µM to select for parasites containing the correct integration. After recovery from drug selection, the population was subcloned via serial dilution and clonal lines were screened for integration of the repair template into the UPRT locus via PCR using P18 and P19. Expression of IF1-HA was confirmed via immunofluorescence and western blotting.

### Quantitative reverse transcription PCR (RT-qPCR)

Total RNA was extracted from lysed tachyzoites using the Zymo Quick-RNA MiniPrep kit (VWR, cat. no. 76020-636). RT-qPCR was conducted using primers P20 and P21 for TgIF1, P22 and P23 for actin, the Luna Universal One-Step RT-qPCR kit (VWR, cat. no. 103307) in an iCycler thermal cycler (Bio-Rad). Relative quantification analysis was conducted using the 2^−ΔΔCt^ method based on actin as a housekeeping gene.

### Western blotting

To prepare samples for western blotting, pellets containing 1 or 1.5×10^7^ parasites were resuspended in 2X Laemmli buffer (20% glycerol, 5% 2-mercaptoethanol, 4% SDS, 0.02% bromophenol blue, 120 mM Tris-HCl pH 6.8) then boiled at 100°C for 5 minutes. Precision Plus Protein Dual Color Standard ladder (Bio-Rad, cat. no. 1610374) was utilized as a molecular weight marker. Following separation by SDS-PAGE, proteins were transferred to 0.2µm pore nitrocellulose membranes (Bio-Rad, cat. no. 1620150) and probed overnight at 4°C on a shaker with mouse anti-Ty (Bastin *et al*., 1996) and rabbit anti-HA (Cell Signaling Technologies, cat. no. 3724S). Membranes were then incubated with goat anti-mouse IgG conjugated to HRP (VWR, cat. no. 102646-160) and goat anti-rabbit IgG conjugated to HRP (VWR, cat. no. 102645-182) secondary antibodies for one hour. Following incubation with enhanced chemiluminescence (ECL) substrate (VWR, cat. no. PI32209), autoradiography film (MTC Bio, cat. no. A8815) was exposed to the membrane and developed via X-ray. After development, membranes were stripped according to manufacturer directions (VWR, cat. no. PI21059) and re-probed with mouse anti-tubulin (Developmental Studies Hybridoma Bank at the University of Iowa, cat. no. 12G10) primary antibody which was used as a loading control. Membranes were then incubated with HRP-conjugated goat anti-mouse IgG secondary antibodies for one hour and again developed using ECL substrate and X-ray film.

### Immunoprecipitation, silver staining and mass spectrometry

Prior to beginning immunoprecipitation, 60µg of mouse anti-Ty antibody (Bastin *et al*., 1996) and 60µg of mouse anti-HA antibody (UGA Bioexpression and Fermentation Facility) were each separately coupled to 1mg of Pierce™ Protein G Magnetic Beads (Thermo Fisher Scientific, cat. no. 88848). Parasites from the IF1^Ty^ and IF1^Over^ strains were lysed at 4°C for 5 minutes in a buffer containing 150 mM NaCl, 20 mM Tris pH 7.6, 1% Triton X-100, 0.1% SDS and supplemented with 1x HALT Protease and Phosphatase Inhibitor (VWR, cat. no. PI78440). Lysates were clarified via centrifugation at 21,000x*g* for 5 minutes at 4°C. The supernatant was incubated with the prepared anti-Ty-coupled Protein G beads (IF1^Ty^) or with the prepared anti-HA-coupled Protein G beads (IF1^Over^) for 1 hour at 4°C. To elute bound proteins, the anti-Ty beads were incubated with 150ng/µl of Ty peptide (Genescript) in lysis buffer and the anti-HA beads were incubated with 150ng/µl of HA peptide (Genescript) in lysis buffer for 30 minutes at 4°C. Elution fractions were resolved by SDS-PAGE then visualized by silver stain as previously described (Shevchenko *et al*., 1996). Precision Plus Protein Dual Color Standard ladder (Bio-Rad, cat. no. 1610374) was utilized as a molecular weight marker. The indicated gel bands were then excised from the gel.

For mass spectrometry analysis, the gel bands were destained with 15 mM potassium ferricyanide and 50 mM sodium thiosulphate solution. After destaining proteins in the gel bands were reduced with 20mM dithiothreitol (Sigma) for 1h at 56^°^C and then alkylated with 60mM iodoacetamide (Sigma) for 1h at 25^°^C in the dark. Proteins were then digested with 12.5ng/uL modified trypsin (Promega) in 50uL 100mM ammonium bicarbonate, pH 8.9 at 25°C overnight. Peptides were extracted by incubating the gel pieces with 50% acetonitrile/5%formic acid then 100mM ammonium bicarbonate, repeated twice followed by incubating the gel pieces with 100% acetonitrile then 100mM ammonium bicarbonate, repeated twice. Each fraction was collected, combined, and reduced to near dryness in a vacuum centrifuge. Samples were then desalted with Pierce Peptide Desalting Spin columns (cat. no. 89852) before running them on the LC-MS.

The tryptic peptides were separated by reverse phase HPLC (Thermo Fisher Scientific Ultimate 3000) using a Thermo Fisher Scientific PepMap RSLC C18 column (2µm tip, 75umx50cm PN# ES903) over a 60-minute gradient before nanoelectrospray using a Orbitrap Exploris 480 mass spectrometer (Thermo Fisher Scientific). Solvent A was 0.1% formic acid in water and solvent B was 0.1% formic acid in acetonitrile. The gradient conditions were 1% B (0-10 min at 300nL/min), 1% B (10-15 min, 300 nL/min to 200 nL/min), 1-3% B (15-15.5 min, 200nL/min), 3-23% B (15.5-35 min, 200nL/min), 23-35 B (35-40.8 min, 200nL/min), 35-80% B (40.8-43.00 min, 200 nL/min), 80% B (43-46 min, 200nL/min), 80-1% B (46-46.1 min, 200nL/min), 1% B (46.1-60 min, 200nL/min).

The mass spectrometer was operated in a data-dependent mode. The parameters for the full scan MS were: resolution of 60,000 across 375-1600 *m/z* and maximum IT 25 ms. The full MS scan was followed by MS/MS for as many precursor ions in a two second cycle with a NCE of 28, dynamic exclusion of 20 s and resolution of 30,000. Raw mass spectral data files (.raw) were searched using Sequest HT in Proteome Discoverer (Thermo Fisher Scientific). Sequest search parameters were: 10 ppm mass tolerance for precursor ions; 0.02 Da for fragment ion mass tolerance; 2 missed cleavages of trypsin; fixed modification were carbamidomethylation of cysteine; variable modifications were methionine oxidation, methionine loss at the N-terminus of the protein, acetylation of the N-terminus of the protein and also Met-loss plus acetylation of the protein N-terminus. Total spectrum count was analyzed in Scaffold; protein threshold was set to 99% and the minimum number of peptides was set to 2.

### Immunofluorescence assays

To confirm mitochondrial localization of IF1^Ty^ and IF1^Over^, as well as to confirm knockout of TgIF1 in the IF1^KO^ strain, co-localization with mitochondrion matrix-targeted SOD2-GFP was utilized. Parasites of each strain were transfected with 20µg of the pT8mycSOD2(SPTP)GFPmycHX plasmid (Pino *et al*., 2007). Immediately following transfection, 40µl of parasites were added to glass coverslips pre-seeded with HFF cells. The next day, intracellular parasites were fixed in a solution of 4% paraformaldehyde for 15 minutes at 4°C. After fixation, a solution of 0.25% Triton X-100 in PBS was used to permeabilize cells for 8 minutes. The coverslips were then blocked for 10 minutes in a solution of PBS containing 5% heat-inactivated fetal bovine serum (IFS) and 5% normal goat serum (NGS). Next, the coverslips infected with IF1^Ty^ or IF1^KO^ parasites were stained with mouse anti-Ty (Bastin *et al*., 1996), while coverslips infected with IF1^Over^ parasites were stained with rabbit anti-HA (Abcam, cat. no. ab9110) primary antibodies for 1 hour. Subsequently, the coverslips infected with IF1^Ty^ or IF1^KO^ parasites were stained with Alexa-647-conjugated goat anti-mouse (Invitrogen, cat. no. A32728), while coverslips infected with IF1^Over^ parasites were stained with Alexa-647-conjugated goat anti-rabbit (Invitrogen, cat. no. A32733) secondary antibodies. Hoechst (Santa Cruz Biotechnology, cat. no. sc-394039) stain was used to visualize nuclei.

Coverslips were mounted onto slides with Prolong Diamond (Thermo Fisher, cat. no. P36961). Images were acquired using an ECHO Revolve microscope and the ECHO Pro application. Image analysis and processing were conducted using Fiji, Adobe Photoshop 2022, and Adobe Illustrator 2022.

### Blue native polyacrylamide gel electrophoresis (BN-PAGE)

To generate samples for BN-PAGE experiments, 2×10^7^ parasites per sample were solubilized in a solution containing 1X NativePAGE sample buffer (Thermo Fisher Scientific, cat. no. BN2008) supplemented with 2.5% digitonin (VWR, cat. no. 10191-280). To create a ladder for accurate estimations of the molecular weight of large membrane-bound complexes, 50µg of bovine heart mitochondria (Abcam, cat. no. ab110338) were solubilized using the same lysis buffer as the parasite samples (Evers *et al*., 2021). Prior to loading into the gel, 1µl of NativePAGE 5% G-250 sample additive (Thermo Fisher Scientific, cat. no. BN2004) was added to each 25µl sample. After separation on a NativePAGE 3-12% Bis Tris protein gel (Thermo Fisher Scientific, cat. no. BN1001BOX), the gel strip containing the bovine heart mitochondria was excised and stained with Coomassie blue (0.3% Thermo Brilliant Blue R-250 (Thermo Fisher Scientific, cat. no. BP101-25), 45% methanol, 10% acetic acid). The rest of the gel containing the parasite proteins was transferred to a PVDF membrane (VWR, cat. no. PI88518). Membranes were probed with rabbit anti-F1β (Agrisera, cat. no. AS05 085) primary antibodies followed by a goat anti-rabbit IgG secondary antibody conjugated to HRP (VWR, cat. no. 102645-182). Following incubation with enhanced chemiluminescence (ECL) substrate (VWR, cat. no. PI32209), autoradiography film (MTC Bio, cat. no. A8815) was exposed to the membrane and developed via X-rays.

### Two-dimensional blue native polyacrylamide gel electrophoresis (2D BN-PAGE)

For 2D BN-PAGE, samples from IF1^Ty^ and IF1^Over^ parasites were prepared and run on the first dimension as described in the previous section for BN-PAGE samples. When the dye front had reached approximately 2/3 of the way down the gel, the electrophoresis was stopped. The lane containing the bovine heart mitochondria ladder was excised and Coomassie stained as previously described. The lanes containing the IF1^Ty^ and IF1^Over^ samples were then carefully excised. Each gel strip was placed in a 1x Laemmli solution (10% glycerol, 2.5% 2-mercaptoethanol, 2% SDS, 0.01% bromophenol blue, 60 mM Tris-HCl pH 6.8) containing 100mM dithiothreitol (DTT; VWR, cat. no. 0281-5G) then microwaved for 10 seconds. Gel strips were then allowed to incubate in the Laemmli/DTT solution at room temperature on a shaker for 5-10 minutes. Following this incubation, each gel strip was carefully loaded horizontally into a 12% polyacrylamide gel poured with a Mini-Protean Prep+1 well comb (Bio-Rad, cat. no.1653367). Precision Plus Protein Dual Color Standard ladder (Bio-Rad, cat. no. 1610374) was utilized as a molecular weight marker for this second dimension run. Samples were run at 120V until the dye front ran off the gel, then were transferred to a PVDF membrane overnight at 25V, 4°C, in transfer buffer containing 0.1% SDS. Membranes were first probed with mouse anti-Ty (Bastin *et al*., 1996) for the IF1^Ty^ sample or rabbit anti-HA (Cell Signaling Technologies, cat. no. 3724S) for the IF1^Over^ sample overnight at 4°C. Membranes were then incubated with goat anti-mouse IgG conjugated to HRP (VWR, cat. no. 102646-160) or goat anti-rabbit IgG conjugated to HRP (VWR, cat. no. 102645-182) secondary antibodies for one hour. Following incubation with enhanced chemiluminescence (ECL) substrate (VWR, cat. no. PI32209), membranes were developed using a BioRad ChemiDoc Imaging System. Following development, membranes were stripped according to manufacturer directions (VWR, cat. no. PI21059) and re-probed with rabbit anti-F1β (Agrisera, cat. no. AS05 085) overnight at 4°C followed by a goat anti-rabbit IgG secondary antibody conjugated to HRP (VWR, cat. no. 102645-182). F1β signal was then captured using the same method.

### Generation of RNA sequencing data

Total RNA was extracted from three biological replicates of lysed IF1^Ty^, IF1^KO^, and IF1^Over^ tachyzoites using the Zymo Quick-RNA MiniPrep kit (VWR, cat. no. 76020-636). The integrity of the extracted RNA was confirmed via an Agilent 2100 Bioanalyzer (Agilent Technologies) using the Eukaryote Total RNA Nano assay. The RNA was then shipped to Psomagen, where the starting total RNA material was quantified via a fluorescence-based quantification method using the Picogreen assay (Thermo Fisher Scientific, cat. no. R11490) on a VictorX2 multilabel plate reader (Perkin Elmer). The RNA integrity was checked using RNA ScreenTape (Agilent Technologies, cat. no. 5067-5576) and RNA ScreenTape Sample Buffer (Agilent Technologies, cat. no. 5067-5577) on a 4200 Tapestation system (Agilent Technologies, cat. no. G2991AA)

500 ng of total RNA served as the input material for library preparation using the TruSeq Stranded mRNA Library Prep kit (Illumina, cat. no. 20020595), along with the IDT for Illumina – TruSeq RNA UD Indexes v2 (Illumina, cat. no. 20040871). The total RNA underwent mRNA purification, involving dilution, the addition of RPB, followed by incubation, and subsequent sequential addition of BWB and ELB, following the specified protocol for the TruSeq Stranded mRNA Library Prep protocol (Illumina, cat. no. 20020595). The isolated mRNA was then fragmented and primed for cDNA synthesis using kit reagents, with an 8-minute incubation at 95°C using a C1000 Touch Thermal Cycler (Bio-Rad, cat. no.185-1196). The cleaved and primed RNA was reverse transcribed into first strand cDNA using SuperScript II Reverse Transcriptase (Thermo Fisher Scientific, cat. no. 18064-014), with Actinomycin D and the FSA (First Strand Synthesis Act D Mix) added to enhance strand specificity. The second strand was synthesized using the 2nd strand master mix from the same TruSeq Stranded mRNA kit, incubated at 16°C for 1 hour. To enable adapter ligation, the double-stranded cDNA (dscDNA) was adenylated at the 3’ end, and RNA adapters were subsequently ligated to the dA-tailed dscDNA. Finally, additional amplification steps were carried out to enrich the library material. The final library was validated using D1000 ScreenTape (Agilent Technologies, cat. no. 5067-5582) and D1000 Reagents (Agilent Technologies, cat. no. 5067-5583) for size information. Quantification was performed using the Quant-iT PicoGreen dsDNA Assay Kit (Thermo Fisher Scientific, cat. no P7589). The validated libraries were then normalized to 10nM and diluted to the final loading concentration of 1.5nM. Utilizing the NovaSeq 1.5 5000/6000 S4 Reagent Kit (300 cycles) (Illumina, cat. no. 20028312) and NovaSeq 1.5 Xp 4-Lane Kit (Illumina, cat. no. 20043131), samples were sequenced on the NovaSeq 6000 system (Illumina).

Analysis of RNA sequencing data

Quality control checks on the IIlumina sequencing data were performed using FASTQC (Andrews, 2010). We then mapped the data to the genome of the *T. gondii* reference strain, ME49 (version 65), using STAR (version 2.7.10b) (Dobin *et al*., 2013). For this purpose, we first generated a genome index using the ‘genomeGenerate’ run mode with the following options set: --genomeSAindexNbases 12, --sjdbOverhang 150, --sjdbGTFfeatureExon exon, --sjdbGTFtagExonParentGeneName gene_id, -- sjdbGTFtagExonParentGeneType gene_ebi_biotype. During the genome indexing step, the reference strain’s genome (FASTA file) and annotations (GTF file) are required. To ensure reads mapping to the untranslated regions (UTRs) of RNA were counted, we modified the GTF file: genomic coordinates with ‘three_prime_UTR’ and ‘five_prime_UTR’ classifications were reassigned to ‘exon’. Next, we mapped the sequencing data using the ‘align reads’ run mode with the following options set: -- alignIntronMin 14, --alignIntronMax 1899, --alignMatesGapMax 497, --quantMode GeneCounts. Finally, we used the featureCounts program in the Subread package (version 2.0.6) to count reads with the following arguments specified: -p -B -- countReadPairs --byReadGroup -s 2 -d 50 -D 600 (Liao *et al*., 2014).

For the differential expression analyses, we used DESeq2 to assess the differential expression (DE) of genes identified in IF1^Ty^, IF1^KO^, and IF1^Over^ parasites (Love *et al*., 2014). Briefly, the output from featureCounts (.txt file) containing count data from all samples was loaded in to R and transformed in to a DESeq2::DESeqDataSet with the design “∼replicate + condition” specified. We discarded genes with less than 9 counts in 3 or more samples prior to running the DESeq2::DEseq() function with the following arguments specified: sfType = “iterate”, fitType = “local”. We extracted the results for IF1^Ty^ versus IF1^KO^ and IF1^Ty^ versus IF1^Over^ comparisons using the DESeq2::results function. We considered genes with a Benjamini and Hochberg adjusted p-value of < 0.05 to be differentially expressed (Benjamini and Hochberg, 1995).

We performed GO Term enrichment analysis on the differentially expressed genes obtained from the DESeq2 analysis using the gprofiler2 package for R (Raudvere *et al*., 2019; Kolberg *et al*., 2020). To this end we use the package’s gost function with the following parameters: organism = “tgondii”, ordered_query = F, multi_query = FALSE, significant = T, exclude_iea = FALSE. measure_underrepresentation = FALSE, evcodes = T, user_threshold = 0.1, correction_method = “fdr”, domain_scope = “known”, custom_bg = NULL, numeric_ns = “”, sources = NULL, as_short_link = FALSE, highlight = F. Gene sets containing 2 or more genes and a fold enrichment FDR value of < 0.1 were considered enriched.

We used the STREME tool (Bailey, 2021) within MEME Suite (v. 5.5.5) to discover enriched amino acid motifs within the subset of genes displaying decreased expression in IF1^KO^ versus IF1^Ty^ and increased expression in IF1^Over^ versus IF1^Ty^. We uploaded a FASTA file containing the amino acid sequences of the proteins for these genes to the MEME suite server. We scanned the amino acid sequences against the PROSITE fixed-length motifs (PROSITE_2021_04) database under the default parameters. The following command was ran on the MEME Suite server: streme --verbosity 1 --oc streme_out -protein --minw 6 --maxw 15 --order 0 --bfile ./background --seed 0 --align center --time 4740 --totallength 4000000 --evalue --thresh 0.05 --p sequences.fa.

### Transmission electron microscopy (TEM)

TEM double fixation and image acquisition were conducted for IF1^Ty^ and IF1^KO^ parasites as previously described (Usey and Huet, 2023). For cristae quantification, 70 sections of each strain that had been pre-selected to contain parasite mitochondria were blinded prior to analysis. Fiji software was utilized to measure mitochondrial area and mitochondrial cristae were counted manually. Following completion of the analysis, images were un-blinded and a student’s t-test was utilized to determine differences in cristae density and the measured mitochondrial area between strains.

### In-gel ATPase assays

In this assay, cellular lysates are prepared and run under native PAGE conditions. Following separation, gels are incubated in a solution containing ATP and lead (II) nitrate. If there is ATPase activity present in the gel, the enzyme will cleave the ATP into ADP and inorganic phosphate, which will then interact with the lead to form a white lead phosphate precipitate on the gel. To prepare the samples for analysis, 5.5×10^7^ or 2×10^8^ parasites from IF1^Ty^, IF1^KO^ and IF1^Over^ strains were solubilized in a buffer consisting of 250mM sucrose, 20mM Tris HCl pH 8, 2mM EDTA pH 8, 750mM amino-N-caproic acid (Sigma Aldrich, cat. no. A-2504-25G), and 2% DDM (Calbiochem, cat. no. 324355). Following clarification of the lysates via centrifugation, 4x NativePAGE sample buffer (Thermo Fisher Scientific, cat. no. BN2008) was added to each sample to a final concentration of 1x. As both a ladder and as a positive control for the assay, 50µg of bovine heart mitochondria (Abcam, cat. no. ab110338) were solubilized using the same lysis buffer as the parasite samples (Evers *et al*., 2021). The samples were separated on a NativePAGE 3-12% Bis Tris protein gel (Thermo Fisher Scientific, cat. no. BN1001BOX) in a 25mM Tris, 190mM glycine running buffer on ice for approximately 3 hours at 150V. The gel was then pre-incubated in a 35mM Tris, 270mM glycine pH 8 buffer for 10 minutes. After this time, the gel was switched to a buffer containing 35 mM Tris, 270 mM glycine, pH 8.4, 14 mM MgCl_2_, 11 mM ATP (Sigma Aldrich, cat. no. A7699-5G), 0.3% (w/v) Pb(NO_3_)_2_ (Sigma Aldrich, cat. no. 228621-100G) (Lacombe *et al*., 2019). The buffer was changed after 4 hours to refresh the reagents and the gel was allowed to incubate in this solution for a total of 20 hours. The gel was then fixed in a 50% methanol solution and lead nitrate precipitates were imaged against a black background. The gel was Coomassie stained for total protein content as a loading control.

### ADP:ATP ratio measurements

ADP:ATP ratio measurements were conducted using the ADP/ATP Ratio Assay Kit (Bioluminescent) (Abcam, cat.no. ab65313) with a protocol adapted from the manufacturer directions. HFF cells were infected with IF1^Ty^, IF1^KO^, or IF1^Over^ parasites. While parasites were still intracellular, the cultures were washed with PBS to remove any extracellular parasites. The intracellular parasites were then syringe released, filtered, and pelleted. The parasite pellets were resuspended in the Nucleotide Releasing Buffer to a concentration of 1×10^7^ parasites/ml and allowed to incubate at room temperature for 5 minutes. During this incubation, the reaction mix was prepared by combining 10% ATP Monitoring Enzyme and 90% Nucleotide Releasing Buffer. 100µl of the reaction mix was to the wells of a white, flat-bottom 96-well plate (Greiner Bio-One, cat. no. 655090) so that the background luminescence levels could be calculated (Data A). 10µl of each lysed parasite solution was then added to wells containing reaction mix and the luminescence was read after 2 minutes (Data B). The luminescent signal was allowed to degrade over 30 minutes before another read was taken (Data C). 10µl of 1x ADP-converting enzyme was added to each well and the luminescence was read after 2 minutes (Data D). The ratio was calculated using the following equation: ADP:ATP ratio = [Data D – Data C]/[Data B – Data A]. Each experiment was conducted in triplicate.

### Cellular ATP concentration measurements

Cellular ATP measurements were conducted as previously described (Usey and Huet, 2023). Syringe released IF1^Ty^, IF1^KO^, and IF1^Over^ parasites were collected in a solution of Fluorobrite DMEM (Thermo Fisher Scientific, cat. no. A1896701) containing 1% IFS and HALT protease inhibitors (VWR, cat. no. PI78440). Pellets were washed in DMEM free from glucose and glutamine (Fisher Scientific, cat. no. A1443001) then resuspended to a final concentration of 6×10^6^ parasites/ml in DMEM free from glucose and glutamine. Aliquots of each parasite sample were immediately flash-frozen to represent initial ATP levels. Additional aliquots of each parasite sample were also incubated for one hour at 37°C and 5% CO_2_ with equal amounts of the following compounds at the listed final concentrations: 5mM 2-deoxyglucose (Sigma Aldrich, cat. no. D6134-5G) + 25mM glucose (Sigma Aldrich, cat. no. G7021-100G) or 5mM 2-deoxyglucose + 2mM glutamine (Sigma Aldrich, cat. no. G8540-100G). After this incubation, these samples were also flash-frozen. To evaluate the ATP levels in each sample, 100µl of CellTiter-Glo reagent (Promega, cat. no. G7572) was added to the wells while samples thawed at room temperature for 1 hour. Luminescence was measured using a Molecular Devices SpectraMax i3x microplate reader. All conditions were conducted in triplicate and ATP levels for each strain were normalized to initial values.

### Mitochondrial membrane potential measurements

Mitochondrial membrane potential measurements were conducted using the lipophilic cationic dye, tetramethylrhodamine ethyl ester perchlorate (TMRE; Sigma Aldrich, cat. no. 87917-25MG). HFFs were infected with IF1^Ty^, IF1^KO^, or IF1^Over^ parasites. While parasites were still intracellular, monolayers were washed with PBS to remove any extracellular parasites. Fluorobrite^TM^ DMEM (Fluorobrite; Thermo Fisher Scientific, cat. no. A1896701) containing 1% IFS was added to each culture before scraping and syringe release of intracellular parasites. Parasites were pelleted then resuspended in Fluorobrite with 1% IFS (unstained), or solutions containing 250nM TMRE, 250nM TMRE with 10µM carbonyl cyanide 4-(trifluoromethoxy)phenylhydrazone (FCCP) (Sigma Aldrich, cat. no. C2920-10MG), or 250nM TMRE with an equivalent amount of DMSO (vehicle control). Parasites were incubated in these solutions for 30 minutes at 37°C and 5% CO_2_. Following staining, parasites were pelleted, and samples were washed then resuspended in either PBS or PBS containing 10µM FCCP or DMSO. Parasites were added in triplicate to a 96 well plate (Greiner, cat. no. 655090) at 2×10^7^ parasites/well. Fluorescence was measured using λ_ex_ 520/25 nm; λ_em_ 590/35 nm filters on a Synergy H1 Hybrid Reader (Biotek).

### Low oxygen plaque assays

To determine the effect of low oxygen conditions on the growth of IF1^Ty^, IF1^KO^, and IF1^Over^ parasites, 500 parasites from each strain were added in triplicate to two 6 well plates pre-seeded with HFFs. One plate containing each strain was incubated under normoxic conditions (21% O_2_, 5% CO_2_, 37°C) while a second plate containing each strain was incubated under hypoxic conditions (0.5% O_2_, 5% CO_2_, 37°C). Plates were left undisturbed in these conditions for 8 days, after which wells were washed with PBS, fixed in 95% ethanol for 10 minutes then stained with a crystal violet solution (2% crystal violet, 0.8% ammonium oxalate, 20% ethanol) for 5 minutes. Wells were subsequently washed again then scanned for analysis. The plaque size of 20 plaques per well and the total number of plaques per well were quantified manually using Fiji.

### Monensin plaque assays

To determine the effect of monensin on the growth of IF1^Ty^, IF1^KO^, and IF1^Over^ parasites, 500 parasites from each strain were added in triplicate to 6 well plates pre-seeded with HFFs. Parasites were allowed to invade the HFFs for 2 hours before the media on one set of wells was supplemented with either 0.003µM monensin (VWR, BUF074) or vehicle control (70% ethanol). After 24 hours of treatment, the wells were washed twice with PBS before fresh media was added. Plates were left undisturbed for 7 days, after which wells were washed with PBS, fixed in 95% ethanol for 10 minutes then stained with a crystal violet solution (2% crystal violet, 0.8% ammonium oxalate, 20% ethanol) for 5 minutes. Wells were subsequently washed again then scanned for analysis. The plaque size of 20 plaques per well and the total number of plaques per well were quantified manually using Fiji.

### Data availability

All raw sequencing data generated in this study can be found in the Sequence Read Archive (SRA) at the NCBI National Library of Medicine (https://www.ncbi.nlm.nih.gov/sra) under the BioProject code: PRJNA1137608. Archived scripts (Shell and R) used to process the RNA sequencing data and its output files as at time of publication are available at Zenodo.

## ABBREVIATIONS

(IF1): ATPase inhibitory factor 1
(DHFR): dihydrofolate reductase
(ROS): reactive oxygen species
(TgIF1): *Toxoplasma gondii* homolog of IF1
(OXPHOS): oxidative phosphorylation

## AUTHOR CONTRIBUTIONS

M.M.U. and D.H. conceived and design the experiments; M.M.U. and A.A.R. performed the experiments; M.M.U., A.A.R. and D.H. analyzed the data. All authors contributed to the article and approved the submitted version.

## ACKNOWLEDGMENTS

We would like to thank the Koch Institute’s Robert A. Swanson (1969) Biotechnology Center for technical support, specifically Richard Schiavoni from the Biopolymers & Proteomics Core for the mass spectrometry data analysis. We would also like to thank Wandy Beatty at the Washington University Molecular Microbiology Imaging Facility for acquisition of our transmission electron micrography images. We appreciate Dr. Chris West and Dr. Msano Mandalasi at the University of Georgia for sharing their equipment and assisting with the hypoxia experiments. We would also express our gratitude to Dr. Silvia Moreno and her laboratory for the feedback, plasmids and antibodies. In particular, we would like to acknowledge Barna Baierna from the Moreno Lab for assistance with the 2D BN-PAGE and native PAGE experiments. Lastly, we would like to thank Kaelynn Parker for experimental assistance throughout the completion of this work.

---

**Figure S1.**
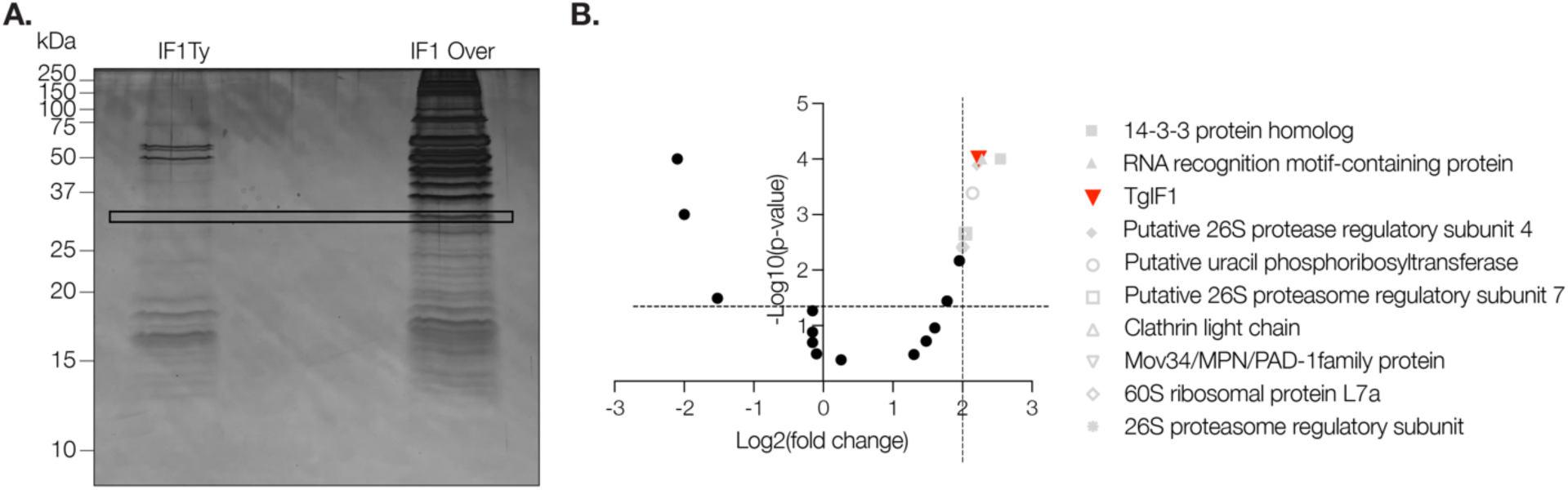
TgIF1 overexpression generates a high molecular weight oligomer. **A**. Following Ty or HA immunoprecipitation, the eluates of IF1^Ty^ and IF1^Over^ parasites were resolved via SDS-PAGE then visualized by silver staining. The indicated bands were excised from the gel then sent for analysis by mass spectrometry. **B** Volcano plot of mass spectrometry results from (**A**). The fold change indicates peptides enriched in the IF1 Over sample compared to the IF1Ty sample. The p-value was generated via Fisher’s exact test. Dotted lines indicate peptides with log_2_(fold change) over 2 and a - log_10_(p-value) over 1.3. Proteins over this threshold, but listed in grey, are contaminants due to their molecular weight being larger than the difference between the low and high molecular weight bands in the IF1^Over^ blot. TgIF1 (red) is highlighted.

**Figure S2.**
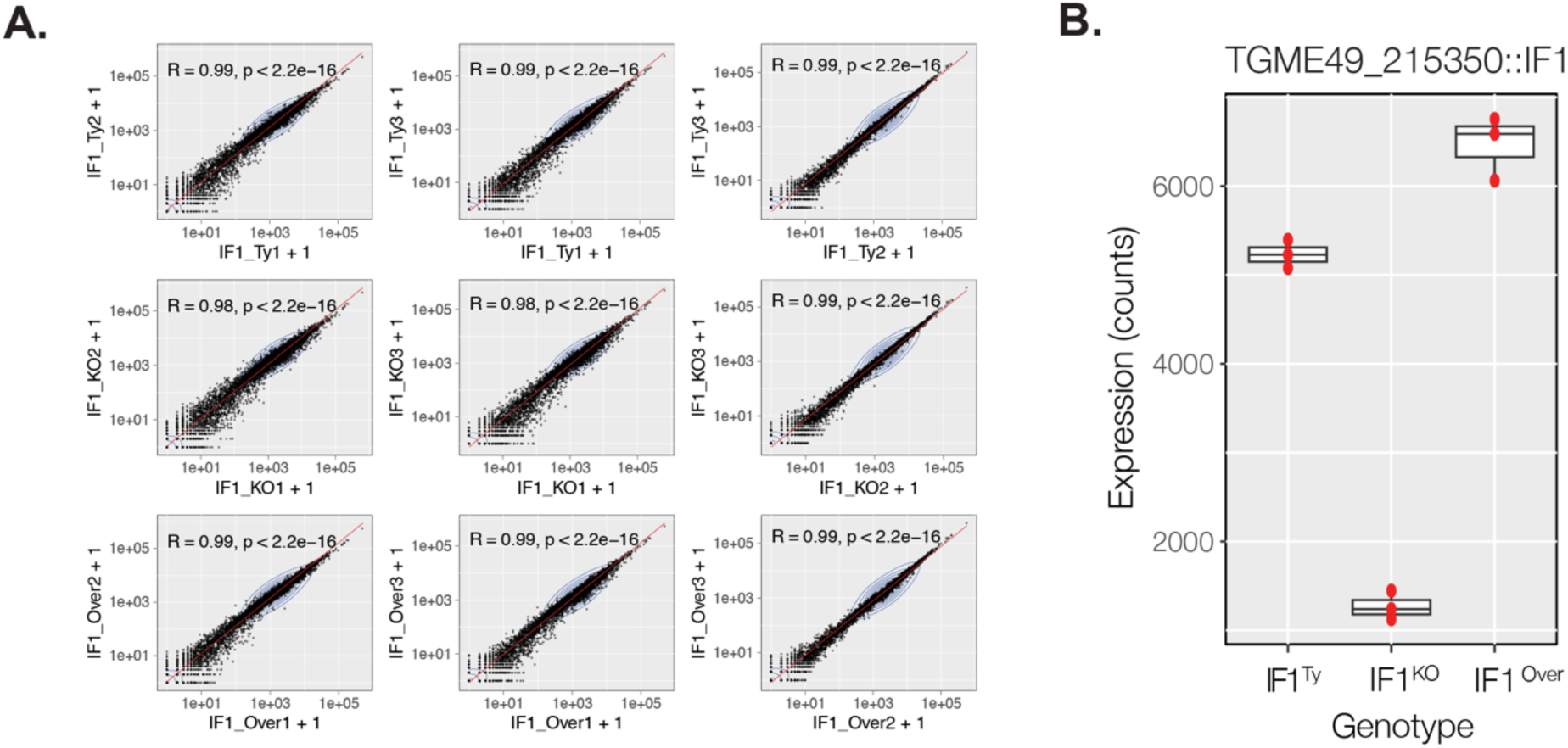
Transcriptomic analysis of parasites lacking and overexpressing TgIF1 reveals altered expression of genes associated with various biological processes. **A.** Scatter plots showing the correlation of gene expression data for each biological replicate processed for RNA sequencing (R = Pearson correlation coefficient). **B**. Boxplots displaying expression levels for TgIF1 in IF1^Ty^, IF1^KO^, and IF1^Over^ parasites. Each point represents expression values obtained from individual RNA-sequencing data sets for each genotype.

